# Rescue of the increased susceptibility to Mild Chronic Oxidative Stress of iNeurons carrying the MAPT Chromosome 17q21.3 H1/H1 risk allele by FDA-approved compounds

**DOI:** 10.1101/2022.11.07.515284

**Authors:** E Sadikoglou, D Domingo-Fernández, N Savytska, N Fernandes, P Rizzu, A Illarionova, T Strauß, SC Schwarz, A Kodamullil, GU Höglinger, A Dhingra, P Heutink

## Abstract

The microtubule associated protein tau (MAPT) chromosome 17q21.31 locus lies within a region of high linkage disequilibrium (LD) conferring two extended haplotypes commonly referred to as H1 and H2. The major haplotype, H1 has been genetically associated with an increased risk for multiple neurodegenerative disorders, including Progressive Supranuclear Palsy (PSP), Corticobasal Degeneration (CBD), *APOE* ε4-negative Alzheimer’s disease (AD) and Parkinson’s disease (PD). The mechanism causing this increased risk is largely unknown. Here, we investigated the role of Mild Chronic Oxidative Stress (MCOS) in neurogenin 2 (*NGN2*) induced neurons (iNeurons) derived from iPS (induced pluripotent stem cells) from carriers of both haplotypes. We identified that iNeurons of the H1 homozygous haplotype showed an increased susceptibility to MCOS compared to homozygous H2 carriers, leading to cell death through ferroptosis. We performed a cellular screen in H1 iNeurons using a FDA-approved Drug Library and identified candidate molecules that rescued the increased susceptibility to MCOS and prevented ferroptosis in H1 iNeurons.

**Highlights:**

- Mild Chronic Oxidative Stress induces neurotoxicity via ferroptosis on iNGN2 neurons
- Axonal degeneration, disordered microtubules, blebs precede neurotoxicity
- MAPT-17q21.3 locus H1/H1, risk allele for NDD is more vulnerable to MCOS
- FDA-approved drugs reverse MCOS induced ferroptosis on H1/H1 risk allele

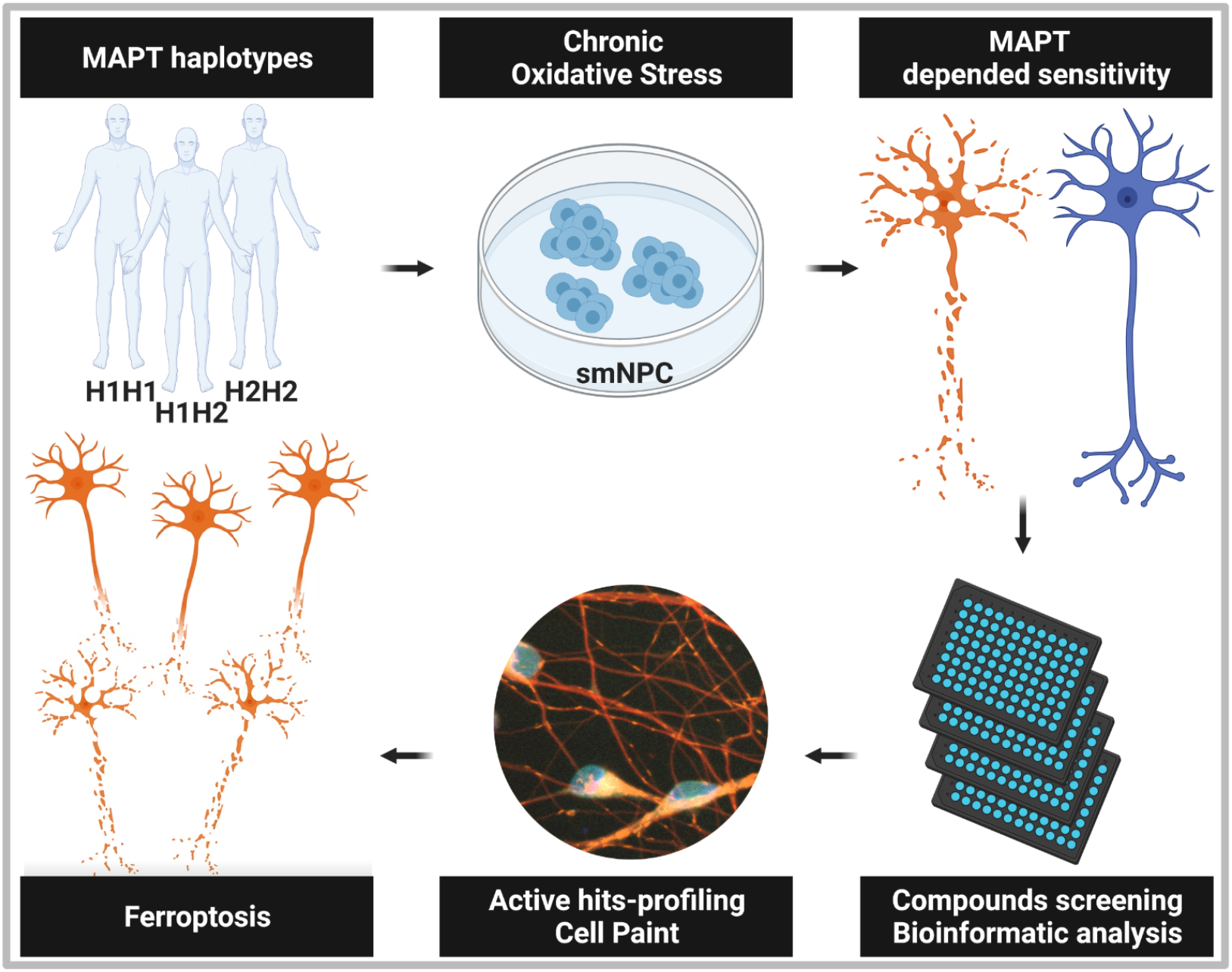

## Introduction

Genome Wide Association Studies (GWAS) have revealed a strong correlation between the Microtubule associated protein tau (*MAPT*) locus on chromosome 17q21.31 and neurodegenerative diseases (NDD), although the causal gene and genomic variants have not yet been identified. The *MAPT* 17q21.31 locus contains a 970-kb inversion and represents one of the most structurally complex and evolutionarily dynamic regions of the genome. This locus occurs in humans as two haplotypes, H1 (direct orientation) and H2 (inverted orientation) which show no recombination between them over a region approximately 1.8 Mb and contains several genes including, *MAPT*, KAT8 regulatory NSL complex subunit 1 (*KANLS1*), Corticotropin releasing hormone receptor 1 (*CRHR1*), Saitohin (*STH*) and N-ethylmaleimide sensitive factor vesicle fusing ATPase (*NSF*) genes (Pittman *et al*., 2004; Caffrey and Wade-Martins 2007; Bowles *et al*., 2022). The major H1 haplotype is considered as the ancestral haplotype while the minor H2 allele is present in 20 % of individuals of European ancestry, is rare in Africans and almost absent in East Asians (Stefansson *et al*., 2005; Hardy *et al*., 2005 ; Campoy *et al*., 2022).

Genetic studies have consistently associated the major H1 haplotype with an increased risk for multiple NDD, including tauopathies such as PSP, CBD and *APOE* ε4-negative AD but also PD (Baker *et al*., 1999; Houlden *et al*., 2001; Pittman *et al*., 2005; Rademakers *et al*., 2005; Zabetian et al., 2007; Höglinger *et al*., 2011; Pastor *et al*., 2016; Sánchez-Juan *et al*., 2019). The H2 haplotype has been subject to recurrent microdeletions associated with the 17q21.31 microdeletion syndrome (Rao *et al*., 2010). Initial studies focused largely on the differential expression of MAPT (Rademakers *et al*., 2005; Caffrey *et al*., 2006; Caffrey *et al*., 2007; Allen *et al*., 2014) and identified the SNPs rs17651213, rs1800547 (Lai *et al*., 2017) and rs242561 (Wang *et al*., 2016) as modulators of MAPT expression, evidence is emerging that other genes at the locus also showed differential expression between the haplotypes (de Jong *et al*., 2012; Bowles *et al*., 2022 Campoy *et al*., 2022). Despite the strong genetic correlation of the MAPT 17q21.31 locus haplotypes with NDD, up until now there is no conclusive functional experimental evidence linking the MAPT 17q21.31 locus haplotypes with distinct molecular consequences leading to neurodegeneration.

The importance of oxidative stress (OS) in the development and the progression of NDD is well documented (Barnham *et al*., 2004; Uttara 2009; Chen *et al*., 2012; Liu *et al*., 2017; Singh *et al*., 2019). Consequences of OS have been extensively studied at molecular level by direct or indirect induction of reactive oxygen species (ROS) in neuronal cultures (Stamer *et al*., 2002; Takeuchi *et al*., 2005; Tang-Schomer *et al*., 2012; Fang *et al*., 2012; Liebrt *et al*., 2016; Datar *et al*., 2019; Yong *et al*., 2019, Fassier *et al*., 2013). Increased ROS levels have been reported to cause axonal degeneration, blebbing and fragmentation due to cytoskeletal disassembly and microtubules disarrangement prior to neuronal death (Whittemore *et al*., 1995, Ricart and Fiszman, 2001; Roediger *et al*., 2003; Fang *et al*., 2012, Barsukova *et al*., 2012, Yong *et al*., 2019). However, the majority of the studies mentioned above were conducted by exogenous treatment with high concentrations agents for short periods of time, on differentiated neurons of animal origins with H1 haplotype (Whittemore *et al*., 1995; Ricart and Fiszman, 2001; Barsukova *et al*., 2012; Fang *et al*., 2012).

In the current study, we examined the effect of MCOS on differentiating neurons in the genetic background of H1 – H2 MAPT-17q21.31 locus haplotypes. We used *NGN2* transduced, small molecule neural precursor cells (smNPCs) derived from nine different iPS lines of healthy donors representing the homozygous H1/H1, H2/H2 and the heterozygous H1/H2 MAPT haplotypes. MCOS in differentiating neurons was induced by removing known antioxidants (AO) like vitamin E, glutathione, superoxide dismutase and catalase from the media of smNPCs and neurons for prolonged time periods. We observed axonal degeneration and microtubules bundles disarrangement prior to cellular death, which is in good agreement with previous oxidative stress (OS) studies (Press and Milbrandt, 2008; Fang *et al*., 2012; Barsukova *et al*., 2012; Datar *et al*., 2019). Axonal degeneration induced by long term ablation of AO was followed by neuronal death via ferroptosis, a relatively recently described iron mediated lipid peroxidation dependent cell death mechanism, strongly associated with the pathophysiology of NDD (Reichert *et al*., 2020). Furthermore, we report here for the first time a significant difference between the MAPT-17q21.31 locus H1 – H2 haplotypes towards their susceptibility to MCOS with the risk allele, H1 haplotype being more sensitive than H2.

In order to identify candidate molecules that could be neuroprotective for the sensitive H1 haplotype against MCOS, we screened a library of FDA-approved molecules. Bioinformatic analysis of already studied and documented primary hits disclosed further implications regarding the cell death mechanism via ferroptosis and the neuroprotection provided by the positive hits. Time lapse imaging and cell painting assay of live neuronal cultures under MCOS with propagating cellular death in neuronal population, lipid peroxidation, plasma membrane rupture, mitochondrial and lysosomal morphology, all characteristic features of ferroptosis, helped to identify the most potent active hits at subcellular level.

## Results

### Distinct response of MAPT-17q21.31 locus haplotypes to mild chronic oxidative stress

To monitor the effect of MCOS on neuronal differentiation, we treated smNPCs of nine different cell lines (three homozygous H1/H1, three homozygous H2/H2 and three heterozygous for H1/H2) in parallel with and without AO for 3, 4 and 6 weeks. iNeurons showed a difference in viability between the H1 and H2 haplotypes after Calcein Red staining (supplementary **Fig.S1A**). H1/H1 neurons were more susceptible throughout the increasing time of AO depletion, while H2/H2 neurons were resistant. The heterozygous H1/H2 showed intermediate sensitivity. Moreover, an additive effect of OS could be observed in all cell lines by increasing time of MCOS (3, 4 and 6 weeks) and /or increasing days of neuronal maturation (day 10, days 12/13). At the smNPC stage we did not observe any differences in conditions with and without AO, in any of the cell lines, even after prolonged periods of treatment **(Fig.S1B - C**).

To assess whether the neuronal viability differences were significantly correlated to the H1 or H2 haplotypes, we normalized the minus AO to plus AO treated neurons, per cell line and the results from two experiments were analyzed with nested one way-ANOVA. The comparison between the mean values of haplotypes are shown in **Fig.1A**. For day 10 neurons, after 3 weeks of treatment without AO, H1/H1 haplotype cell lines were significantly more sensitive than H2/H2 (p<0.001). The difference becomes less significant with increasing time of treatment (p<0.0026 for four weeks and p<0.0261 for six weeks), since some neurotoxicity was observed in H2/H2 lines as well. In more mature neurons (days 12/13), the differences in the mean values per haplotype were not significant, as some of the H2/H2 lines were affected also from AO depletion.

**Fig. 1:**
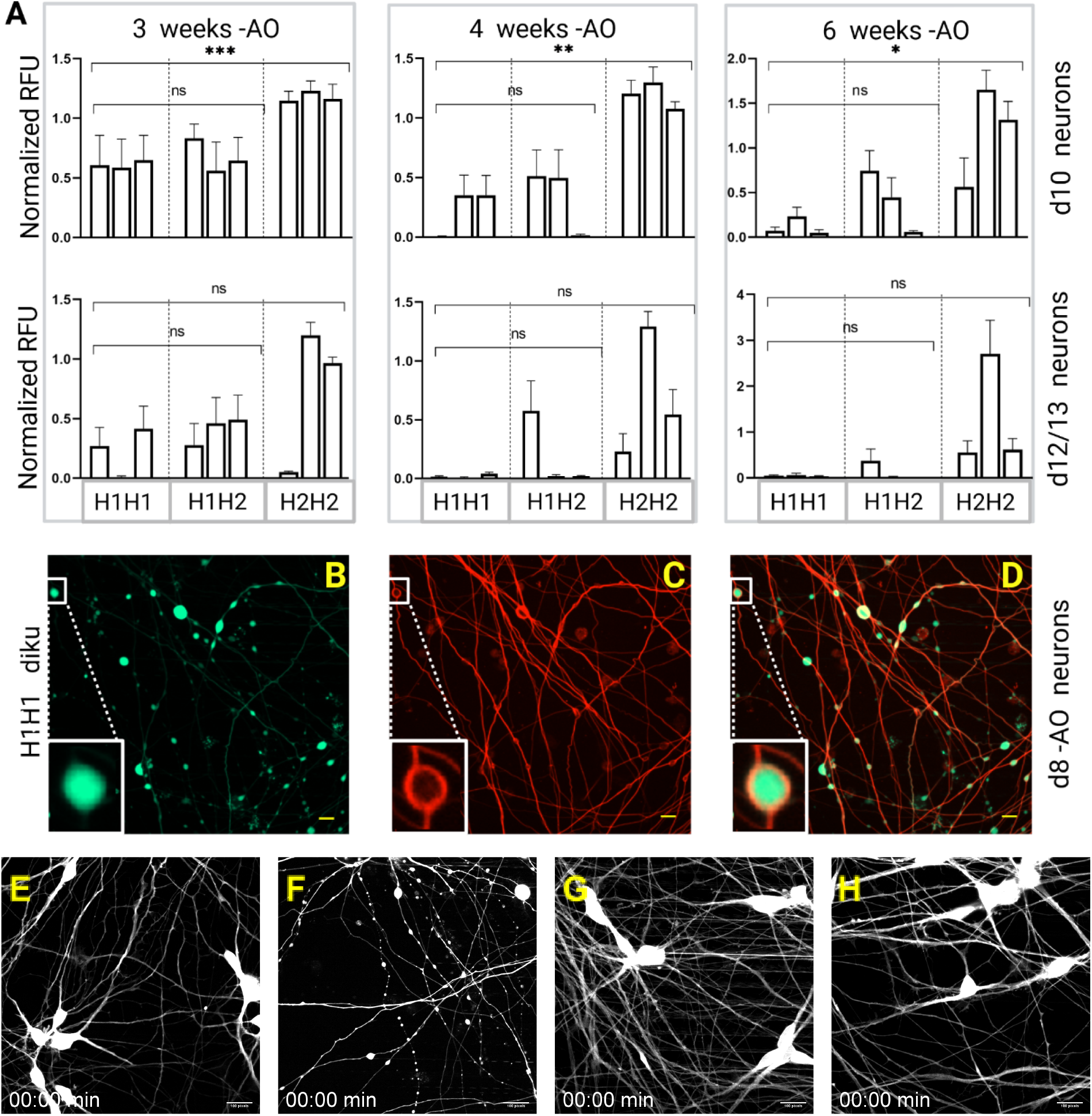
Difference of MAPT haplotypes to mild oxidative stress. **(A)** Neuronal viability microlate assay. Relative fluorescence units from Calcein staining. Three cell lines per haplotype treated without AO were normalized to treated with AO cells and analyzed with nested one-way ANOVA. Dunnett’s test was used for the correction of values in multiple comparisons of haplotype groups. Data represent mean values ± SFM from two experiments with three biological replicates and P< 0.05: *, P< 0.01 : **, P< 0.001: **, P<.0001 : ****. **(B-C-D)** Fluorescence imaging of live neuronal staining with Calcein Green **B** and microtubule tracker C. Blebs and microtubule bundles disarranged regions overlap in **D**, scale bar 10μm. **(E-F-G-H)** Time lapse for 5 hours imaged every 20 minh of day 8 neurons stained with Calcein from sinNPC’s treated for 5 weeks with and without AO. E H1/H1 diku with, **F** without, and **G** J12/112 uilk with and **H** without AO. Scale bar 100 pixels on ImageJ For complete time-lapse videos refer to **Videos S1-4** respectively.

To exclude any possible technical effects, we repeated the experiment from the iPS stage, generated novel smNPCs and challenged one H1/H1 line (diku) and two H2/H2 lines (zihe and uilk) as before with MCOS. We confirmed that the H1/H1 neurons were more sensitive to AO depletion than H2/H2 after five weeks of treatment (data not shown).

The observed neurotoxicity was due to AO depletion, because supplementation of media with AO at day 5 of neuronal differentiation was sufficient to rescue the cells from neurotoxicity at the levels of neurons treated with AO, for all the cell lines (data not shown). A detailed time course of rescuing the neurotoxicity with AO supplementation was done also for the screening assay set up (see material and methods and **Fig.S2**)

### Axonal degeneration and Ferroptotis due to mild chronic oxidative stress

During the course of the neurotoxicity assay, we noticed increased signs of axonal degeneration similar to those observed in tauopathies before cell death (Kneynsberg *et al*., 2017). Blebs, also referred to in the literature as swellings, beadings, spheroids or varicosities, (Luo and O’Leary, 2005; Datar *et al*., 2019; Yong *et al*., 2019; Palumbo *et al*., 2021) were observed at day 8 neuronal axons of H1/H1 haplotype, depleted for 5 weeks of AO, just before the neuronal cell death. These blebs started to develop at sub-regions of wells and propagated to other regions within 24 hours. Towards the end of this time frame the neuronal cell body adapted to a more round and swollen shape, just before their death (**Fig.S1E**). The H2/H2 haplotype cell lines treated in parallel without AO (**Fig.S1G**), and the AO treated H1/H1 cell lines (**Fig.S1D** and **F**) retained intact axonal networks throughout the differentiation. In a series of time lapse images of H1/H1 and H2/H2 neurons differentiated from smNPCs, treated with (**Fig.1E** and **1G** respectively) or without AO (**Fig.1F** and **1H** respectively) for 5 weeks, we studied the spatiotemporal progression of axonal degeneration until the neuronal death on day 8 neurons. The formation of blebs in H1/H1 neurons treated without AO started as small elongated regions on axons, gradually increased in size and numbers and axons thinned progressively before the neuronal death (**Fig.1F**). Evidence of the above-mentioned cascade of events, other than some reversible swellings due to big cargo transport movements, were absent in H2/H2 lines treated without AO (**Fig.1H**). In contrast, we did not observe any differences in the axonal network of all cell lines of both haplotypes treated with AO (**Fig.1E** and **1G** respectively).

Previously, it has been shown that blebs during axonal degeneration, due to OS induced by different perturbations on neurons, indicate regions of disordered microtubules bundles (Fassier *et al*., 2013; Datar *et al*., 2019). Staining with Calcein (**Fig.1B**) and tubulin tracker (**Fig.1C**) of day 8 neurons of diku (H1/H1), treated for 5 weeks without AO, confirmed that the blebs indeed overlap with regions of disorganized microtubules bundles (**Fig.1D**).

An intriguing observation was related to the mode of cellular death in the neuronal culture, which differed from other forms of regulated cell deaths (Riegman *et al*., 2019). The signs of axonal degeneration and subsequent cell death were spreading and propagating between cells in a wave-like manner which was shown earlier to be characteristic of ferroptosis, a recently described form of programmed cell death characterized mainly by implication of iron and lipid peroxidation (Davidson and Wood, 2020; Riegman *et al*., 2020; Nishizawa *et al*., 2021). We therefore treated cell lines (three per haplotype) without AO for 5 weeks and stained them with the LIVE/DEAD kit on days 9, 12 and 21 of neuronal differentiation. In a series of time lapse images for 24h of 3×3 tiled fields and several regions within the wells, we observed the propagation of axonal degeneration and neuronal death in a wave like manner in all cell lines of both haplotypes (**Fig.S1H** to **M**). The difference between the haplotypes was the day of neuronal differentiation at which the axonal degeneration and the subsequent neuronal death initiated, as it was on average earlier for H1/H1 than in H2/H2 cell lines in agreement with our previous cytotoxicity assay (**Fig.1A** and **Fig.S1A**).

### Small molecules library screening

After elucidating the difference in response to AO depletion between the H1 and H2 haplotypes, we developed a screening assay (see material and methods and **Fig.S2**) to identify candidate molecules that could rescue and reverse the detrimental effect of MCOS on H1/H1 neurons. For the screening assay, we used a library of FDA-approved drugs (FDA-approved Drug Library, Selleckchem), with well-studied mechanisms of action of the comprised bioactive compounds. The screening protocol (**Fig.2A**) included neurons kept constantly in without AO medium as negative control, whereas positive control neurons were rescued on day 6 of neuronal differentiation with medium supplemented with AO. The screen was conducted in two independent batches (**Fig.2B** and **2C**) and the assay quality was determined by the z-factor which was on average 0.87 for the first batch and 0.68 for the second batch **(Fig.S2D**).

**Fig. 2:**
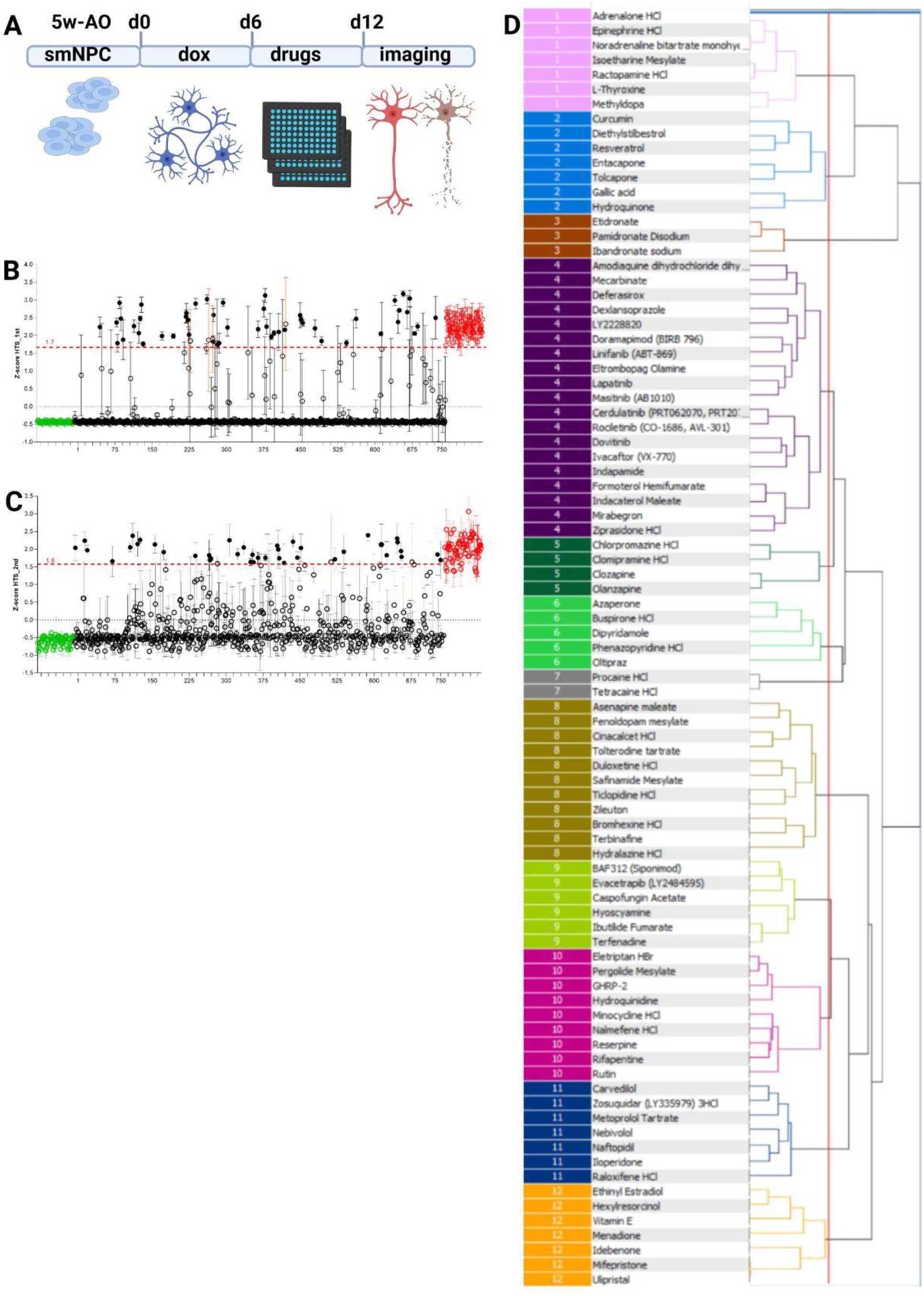
FDA approved drugs library screening. **(A)** Schema of the screening assay. Cortical neurons were differentiated from smNPC’s treated without AO tor 5 weeks. After standard medium change on day 3, compounds were added on day 6 along with medium change. Cell viability was assessed by imaging on day 12 with calcein staining. Screening was completed in two independent runs with n=4 replicates. **(B)** and **(C)** z-scores of compounds calculated with HitSeekR, are shown from first and second runs. Green and red points are the negative and the positive controls respectively. Compounds are shown with black dots the ones above and black circles the ones below the cut off (red dotted line), Hits with SD>0,7 from n=4 replicates (orange error bars) were excluded, **(D)** Structure similarity analysis of primary hits. Exact SAHN clustering dendogram of hits based on their Day Light fingerprints and Ward’s linkage methods with Scaffold Hunter 2.6.3. (Schafer *et al*., 2017).

Live neuronal counts were analyzed with the web application HitSeekR (List *et al*., 2016) and hits were defined as the small molecules that had z-scores >1.7xSD of positive control in the first and >1.6xSD in the second batch (**Fig.2B** and **2C**). 6.7% of the initial library was determined as primary hits, including several small molecules with already known AO activity. The inclusion of Deferasirox, Dexlansoprazol, Idebenone, Zileuton, Vitamine E, Ethinyl-Estradiol, Lapatinib, Carvedilol and Curcumin, well-documented ferroptosis inhibitors in earlier studies (Stockwell *et al*., 2017; Mou *et al*., 2019; Tang *et al*. 2021) among our primary hits confirmed our previous observations of cell death mechanism. On the other hand, the pan-caspase - Apoptosis inhibitor Emricasan, (Barreyro *et al*., 2015) and quite a number of autophagy inhibitors (Omeprazole, Nimodipine, Azithromycine, Chloroquine, Hydroxychloroquine) (Nakahira *et al*., 2013; Draf *et al*., 2021), which were part of our initial library, could not rescue the MCOS induced neurotoxicity.

### Structure similarity analysis of active hits

Substructure similarity analysis of drugs has been used successfully in the past for selection and ranking of hits after high throughput screening assays (Martin *et al*., 2002). Additionally, it can be the groundwork of structural analogy studies, for the design and the development of new drugs (Kafarski and Lipok, 2015).

Clustering of our primary hits with sequential agglomerative hierarchical non-overlapping (SAHN) method based on DayLight fingerprints of compounds and Ward’s Linkage method (Scaffold Hunter 2.6.3 Schäfer *et al*., 2017) returned 12 groups of structurally similar small molecules (**Fig.2D**). Setting up the threshold lower to obtain more clusters was not necessary as it would separate the Gallic acid and the Hydroquinone from Cluster 2 into an independent cluster (**Fig.2D** and **Table S1** Cluster 2 blue dashed compounds). On the other hand, setting the threshold higher to obtain less clusters would merge Clusters 9 and 10 to a more structurally diverse and complex group (**Fig.2D** and **Table S2**).

The substructure similarity analysis in some cases, especially in small clusters like 3, 5, and 7 (**Fig.2D, Tables S1** and **S2**), also gave functionally related drugs with similar therapeutic uses per cluster. Cluster 3 differed quite early by the rest as it was consisted of bisphosphonates with a phosphorus-carbon-phosphorus structure (Allgrove, 1997) and are used in the management of calcium and bone metabolism (Vasikaran, 2001; Roelofs *et al*., 2010) (**Table S1**).

Various known natural AO were grouped in Clusters 2 and 12 (**Tables S1** and **S2**). The first two clusters consisted mostly of polyphenolic compounds like phenylalanine derivatives and catecholamines in Cluster 1 and stilbenes and flavonoids with known antioxidant activities in Cluster 2. Cluster 4 which was the biggest and the most diverse group had antineoplastic and anti-inflammatory drugs. All the tyrosine kinase inhibitors, together with quinoles, and indoles consisted mostly of the group. The tricyclic antidepressants (TCA) composed very early a distinct group in Cluster 5. (Detailed clustering of primary compounds can be found in supplementary **Tables S1** and **S2**).

### Pathway enrichment analysis

Substructure similarity analysis of primary hits and the molecular targets, provided by Selleckchem (**Table S1** and **S2**), demonstrated a wide range of chemical structures and molecular activities. In order to gain further insights on the pathways targeted by the primary hits, we performed pathway enrichment analysis by extracting chemical-protein interactions from DrugBank (Wishart *et al*., 2008) and from the scientific literature (SCAIView; https://academia.scaiview.com). The complementarity of both approaches enabled us to not only rely on well-studied interactions, but also leverage other interactions mined from the literature (**Fig.3A**, step 1).

**Fig. 3:**
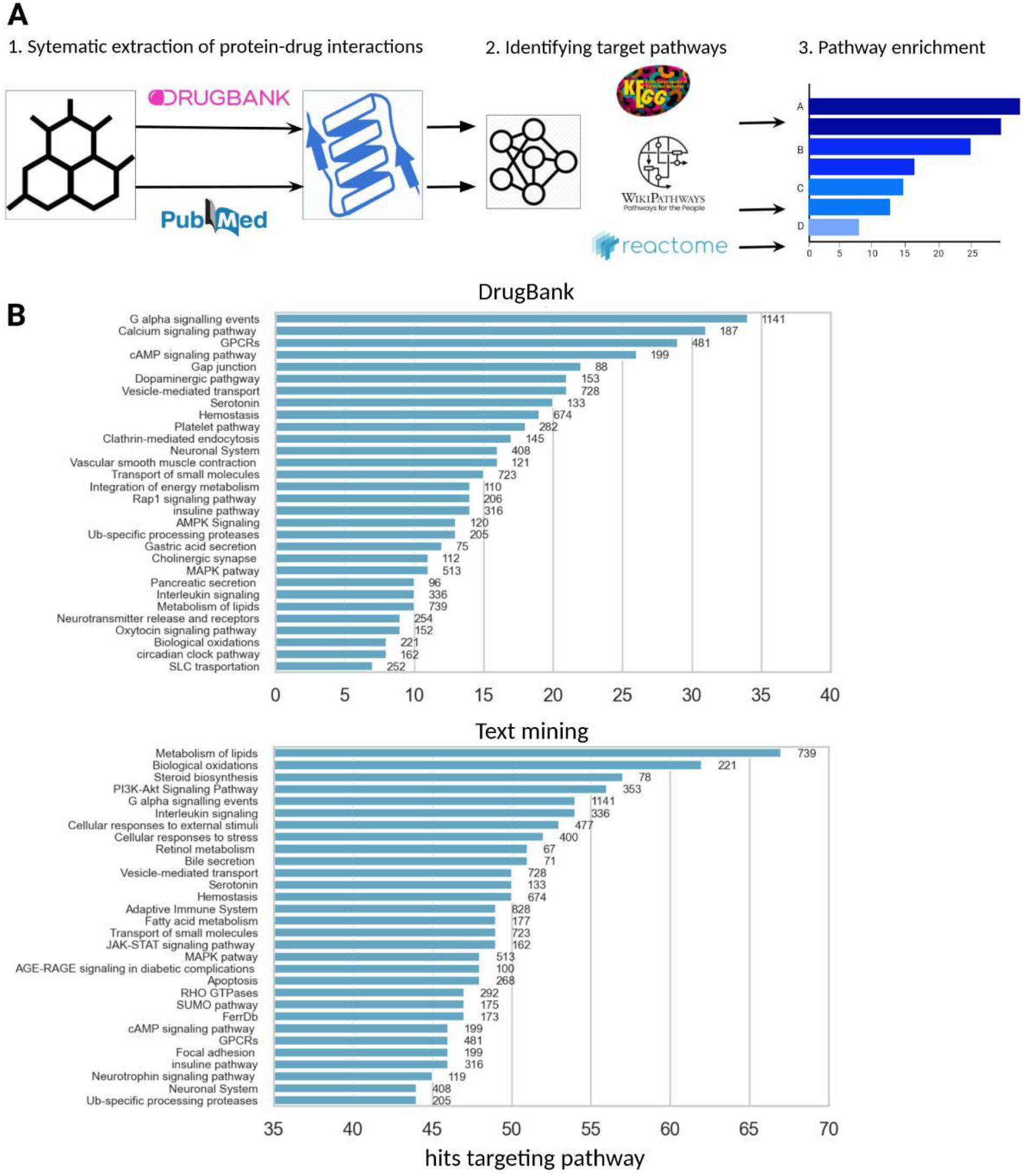
Pathway enrichment analysis of primary hits. **(A)** Bioinformatics analysis scheme employing two complementary approaches. Primary hits target genes obtained from DrugBank and with the text mining of studies in literature. Both sets of gene targets were mapped to pathways in three major pathway databases KEGG, Reactome and WikiPathways and grouped together using their pathway hierarchy. **(B)** Comparison of the enriched pathways targeted by primary hits with both approaches, The Y-axes display the 30 most enriched pathways, the X-axes the number of primary hits targeting each pathway.

As expected, the text mining approach yielded a larger number of proteins being targeted by each primary hit. Interestingly, the targeted pathways and their ranking based on the number of small molecules that targeted them varied significantly between the two approaches. In the case of the DrugBank, the most targeted pathways were: G alpha signaling events, Calcium signaling pathways and G-Protein Coupled Receptors, with 30 primary hits targeting each of these pathways on average (**Fig.3B**). Whereas, in the text mining approach, the most targeted pathways were the lipid metabolism and the biological oxidation pathways, with more than 60 small molecules out of 87 primary hits targeting each of these pathways (**Fig.3B**).

We also used the gene set curated from FerrDb (Zhou and Bao, 2020), as none of the three databases, KEGG (Kanehisa and Goto, 2000), Reactome (Jassal *et al*., 2020) and WikiPathways (Martens *et al*., 2021) had integrated a ferroptosis related pathway. Based on the DrugBank interactions, the primary hits were not related to any of the cell death pathways, as opposed to the text mining interactions where both the ferroptosis and the apoptosis pathways were targeted by more than the half of our primary hits, with 47 and 48 small molecules, respectively. Although the two cell death pathways looked to be equally enriched, when we normalized by the pathways size (i.e., 173 genes in the ferroptosis and 268 in the apoptosis pathway), ferroptosis was more enriched as 27% of the total genes in this pathway were targeted by our primary hits versus 18% of the apoptosis pathway.

In parallel, we checked the number of our primary hits that have already been investigated in the context of neurodegenerative disorders, querying ClinicalTrials.gov using identifiers of the 94 diseases belonging to the MeSH branch of “Neurodegenerative Diseases” (date of the query 13.05.2021). The resulting query (data not shown) revealed that 34 small molecules of primary hits have already been investigated in a clinical trial for any neurodegenerative disease.

In summary, based on the molecular targets provided by Selleckchem and the pathway enrichment analysis, with chemical-protein interactions obtained from DrugBank, primary hits were not closely related to OS and ferroptosis. However, the more inclusive and up to date text mining results strongly supports the neuro-protective effects of primary hits in the observed MCOS induced ferroptosis. This discrepancy is likely due to one of the limitations of databases, which fail to cover the constantly accumulating most actual information presented in scientific articles. Lastly, we would like to note that the text mining workflows were crucial to augment pathway enrichment analysis.

### Dose Response Curves of primary Hits

To gain more insights into the potency and efficacy of all primary hits in reversing the neurotoxicity of MCOS, we performed a Dose Response Curve (DRC) analysis.

Plate reformatting of 87 primary hits was done manually and 5 concentrations of fivefold dilutions (from 5 μ? to 8 nM) of each small molecule was tested in n=4 replicates. Live neuronal numbers were fitted to dose-response curves versus the logarithmic scale of small molecules concentrations (x axes), plotted in groups based on their chemical fingerprint clustering. Results showed a wide variety of DRC (**Fig.S3**) which were classified based on their asymptotes, their fit to the data (r^2^) and their efficacies as in previous studies (Inglese *et al*., 2006; Ribbens *et al*., 2012). Detailed examination of classified hits (**Fig4.A**) showed that we did not obtain any false positive hits from our primary screening assay, since all 87 small molecules were still active at 5 μM. However, we did observe and excluded from further analysis 23 hits that were classified as single dose active small molecules since they rescued the neurotoxicity only at the highest concentration of 5 μM. Almost half of our primary hits (40) showed complete curves with both asymptotes and were further sub-classified based on their efficacy and their fit. Nineteen of the primary hits had partial curves and 2 of them were in the subgroup of partial efficacy with good fit and were excluded also from further validation. Finally, we obtained 5 small molecules which were active at the four highest concentrations of DRC. One of them was Resveratrol, a very well-known AO with multiple neuroprotective activities (Singh *et al*., 2013) and numerous targets in the antioxidant pathway like, free radicals, lipid peroxidation, enzymatic activity and gene expression regulation (Gu *et al*., 2021).

**Fig. 4:**
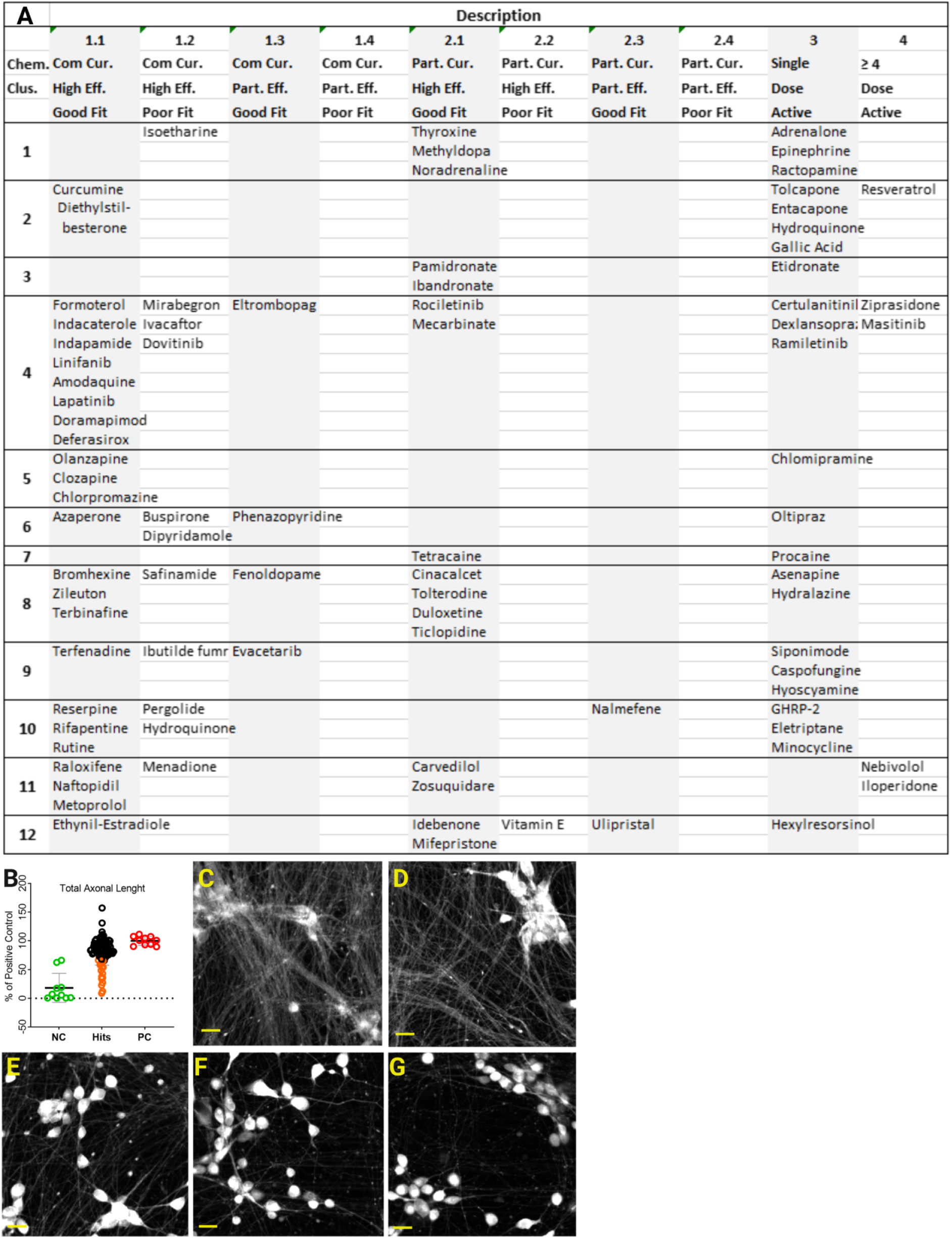
Description of primary hits based on dose response curve (DRC) and total axonal lengh (TAL) analysis. **(.A)** Primary hits classitified based on their DRC asymptotes. their fit to the data(r^2^) and their efficacies. **(B)** TAL analysis of primary hits (black and orange circles) expressed as % of positive control (red circles). With orange circles primary hits scored 2XSD lower than positive controls. Negative controls fire indicated with green. Each data point is the mean of n=4 replicates. **(C-D)** Reprentaitive fileds of pnositive control wiith intact axonal network. **(E-F-G)** Carvedilol, Caspofungin Acetate and Hyoscyamine, compounds with the lower % of positive control in TAL, analysis (orange circles in B). Scale bar 20 μm.

### Total Axonal Length Analysis

Since we observed that axonal degeneration precedes neuronal death in our MCOS model, we investigated if any of our primary hits could have retained the live neuronal body count but still showed degenerated axons. Accordingly, we performed a retrospective total axonal length (TAL) analysis by using the maximum projection of two z-stacks images from the 5 μM concentration of the DRC experiment. In **Fig.4B** the percentage of total axonal length of each small molecule versus the positive control can be seen. In total 19 small molecules (orange circles) of our primary hits (black circles) scored two standard deviations lower than the average of positive control (red circles) and were excluded from further validation. Among them we could not observe any enrichment of a specific chemical cluster, whereas six of them were excluded earlier as single dose active molecules in DRC analysis. In comparison of representative fields of positive control (**Fig.4C** and **D**) versus Carvedilol (**Fig.4E**), Caspofungin Acetate (**Fig.4F**) and Hyoscyamine (**Fig.4G**), small molecules which failed in TAL analysis, the differences in axonal network were obvious.

### Validation of Hits with Cell Painting Assay

Primary and secondary screens revealed 49 FDA-approved compounds which could preserve an intact axonal network and reverse the neurotoxicity due to ferroptosis in a dose dependent manner. As it is known, OS causes damage to macromolecules such as nucleic acids, proteins and lipids (Lassmann and van Horssen, 2016), thus we investigated how healthy neurons were, at subcellular level, after being rescued by these small molecules. For this, we followed a Cell Painting approach (Gustafsdottir *et al*., 2013; Rohban *et al*., 2017) by labeling live neuronal compartments with fluorescent bioprobes to quantitatively profile multiple parameters specifically related to OS and ferroptosis phenotypes.

The aim of the Cell Painting assay was to monitor live neurons under MCOS during the dying process in real time. Images of plasma membrane, lipid peroxides, nucleus, mitochondria, lysosomes and microtubules were taken on days 8 and 9 of neuronal differentiation according to the cascade of events induced by MCOS. Image analysis was performed with CellPathfinder and the extracted features after quality control within and among replicate plates and the removal of all the total not averaged values, approximately 400 features were used in dimensionality reduction with Principal Component Analysis (PCA).

Results showed that the first five and four principal component scores (PC scores) for days 8 and 9 of neuronal differentiation respectively (**Fig.5B** scree plots) were explaining on average the 75% of the variation in our data set. Graphs of variables revealed an absolute clustering of extracted features based on the staining and the phenotype, with the correlation in between them as expected. For example on day 8 (**Fig.5C**) we observed a tight cluster related to the lipids and the oxidized lipids features (green circle) as the most contributing variables to the PC score 1 (denoted as Dim1). The dead nucleus related features cluster (dark blue circle) was in negative correlation with live nucleus features cluster (light blue circle) and the most contributing factors to PC score 2 (denoted as Dim2). As for the day 9 variables graph (**Fig.5D)**, the live nucleus related features cluster (light blue circle) was contributing the most to the PC score 1 and was negatively related with the dead nucleus cluster (dark blue circle) on PC score 2 (denoted as Dim1 and Dim2 respectively). An unexpected result was that the PC score3 which was defined by the lysosomal related features (orange circle). Image comparison between positive and negative controls revealed a clear difference in numbers and sizes of lysosomes within the cells and on the axons (**Fig.5A**). Although the exact mechanism is not yet so clear, the implication of lysosomes in ferroptosis is supported by previous studies from the literature (Torii *et al*.,. 2016; Gao *et al*., 2018; Tian *et al*., 2021; Wang *et al*., 2021).

By using the first 5 PC scores we 3d plotted the active hits as PC1-2-3 (left) and PC1-4-5 (right) for day 8 (upper panel) and for day 9 (lower panel) of imaging (**Fig. 5E**). With the help of 90% of ellipses drawn based on the Mahalanobis distance of individuals from the centroids of the groups we could exclude further nine drugs from the active hits list. Representative fields of positive and negative controls (left and middle columns) and extreme cases of active hits which were excluded by PCA analysis (right column) with the corresponding fluorescence probes per lane are shown in **Fig. 5A**.

**Fig. 5:**
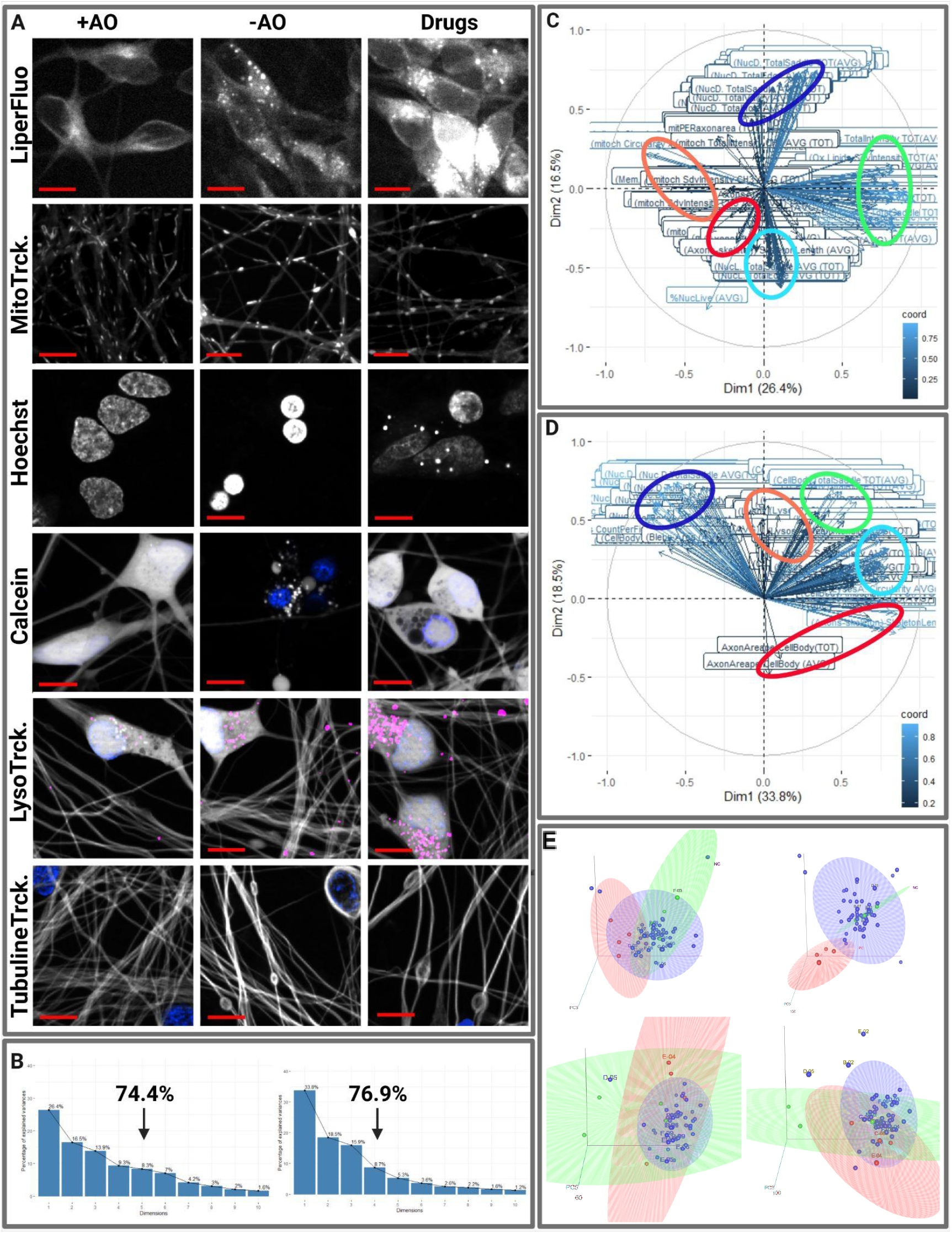
Cell Painting assay. **(A)** Representative fields of neurons treated with (left column) and without AO (middle column). Extreme cases of active hits (right column). Labelled per lanes. the Fluorescence probes used for imaging. Scale bar 10 μM. **(B-C-D-E)** Principal Component Anaysis (PCA) of-cell painting assay. **(B)** Scree plots with the cumulative% of variation in data indicated with arrows for day 8 on the left, and day 9 on the right **(C-D)** Day 8 and day 9 variables graphs with contributions to the first two principal components (Dim 1 and Dim2). **(E)** 3d animating plots of principle components PC1-2-3 (left) PC1-4-5 (righl) day 8 upper panel and day 9 lower panel. 90% ellipses drawn based on the Mahalanobis distance from the centroid of positive controls. (red), negative control (green) and active hit (blue). For complete animating 3d plots refer to **Videos S5-8**.

## Discussion

GWAS have consistently identified the MAPT-17q21.31 locus (H1 – H2) haplotypes as a common risk factor for NDD, including tauopathies such as PSP, CBD and AD (Rademakers *et al*., 2005; Zabetian *et al*., 2007; Höglinger *et al*., 2011; Sánchez-Juan *et al*., 2019). Although the genetic architecture of the locus has been studied extensively, little is known about the molecular mechanism and the causal variant(s) leading to the increased risk associated with the H1 haplotype (Pittman *et al* 2005; Oliveira *et al*., 2004; Rademakers *et al*., 2005; Caffrey *et al*., 2006; Caffrey *et al*., 2007; Allen *et al*., 2014; de Jong *et al*., 2012; Bowles *et al*., 2022 Campoy *et al*., 2022). Therefore, we performed a combinatory study of genetic variation within the context of cellular oxidative stress, another major risk factor in many NDD (Andersen, 2004; Shukla *et al*., 2011; Salim, 2017). Instead of exogenous applications of high concentration of chemicals, as is common in the field, we instead induced MCOS conditions by removing known AO for prolonged time periods from smNPCs and neurons, as a more relevant model to the pathophysiology of NDD.

ROS with pleiotropic effects in CNS, are crucial for the proliferation, maintenance and the lineage specification of mammalian neural stem and progenitor cells (Iqbal and Eftekharpour, 2017; Hameed *et al*., 2015; Nugud, 2018). On the other hand, ROS are neurotoxic with vulnerability depending on the neuronal type and the maturation stages (Wang and Michaelis, 2010; Satish Bollimpelli and Kondapi, 2015). Our results after depleting smNPC of both H1 and H2 haplotypes of AOs and differentiating into iNGN2 cortical neurons point in the same direction. We observed that under MCOS conditions the smNPCs proliferation and the initial stages of neuronal differentiation proceeded normally until the stage where the axons started to form a complex network. The subsequent neurotoxicity, preceded by signs of axonal degeneration, was increased by longer periods of AO depletion and/or more advanced neuronal maturation. Supplementation with AO during the neuronal differentiation prevented the neurotoxicity in a dose dependent manner.

An unanticipated finding was the significant difference in resistance to AO depletion between the H1 – H2 haplotypes which has not been observed before and might have been missed or undetected under the harsher OS conditions that are often used. By comparing three cell lines per haplotype from healthy donors, we demonstrated that the H1/H1 homozygotes where more sensitive to MCOS than the H2/H2 homozygous lines as on average they developed signs of axonal degeneration and neurotoxicity at earlier days of neuronal maturation. To our knowledge, this is the first study establishing a clear phenotypic difference between iNeurons carrying different MAPT-17q21.31 locus haplotypes against the deleterious effects of OS. We hypothesize that our observation could at least partly shed light on the molecular mechanisms leading to the H1/H1 haplotype being a common risk factor for NDD.

Axonal degeneration, a pathological hallmark of numerous NDD (Stokin *et al*., 2005; Clark Jayden *et al*., 2016; Salvadores *et al*., 2017; Chen *et al*., 2011), including tauopathies (Kneynsberg *et al*., 2017), occurs before the neuronal cell death and even before the onset of clinical symptoms (Stokin *et al*., 2005; Adalbert *et al*., 2013; Tagliaferro *et al*., 2016). At the molecular level it has been shown that the directly or indirectly induced ROS levels causing axonal degeneration, blebbing/swelling and fragmentation are due to disarrangement of microtubules bundles prior to neuronal death (Whittemore *et al*., 1995, Ricart and Fiszman, 2001; Roediger and Armati, 2003; Fang *et al*., 2012, Barsukova *et al*., 2012, Datar *et al*., 2016; Yong *et al*., 2019). By live neuronal staining and time-lapse imaging, we indeed observed axonal degeneration and microtubule bundles disarrangements before the neuronal death in H1/H1 cell lines, as a result of MCOS, which appeared at later time points in H2/H2 lines treated in parallel.

Our observation that during the neurotoxicity assay axonal degeneration and subsequent neuronal death, in all cell lines of both haplotypes, was spreading and propagating between neurons in a wave-like manner suggested that ferroptosis could be the mechanism for the observed cell death (Dixon *et al*., 2012; Davidson and Wood, 2020; Riegman *et al*., 2020; Nishizawa *et al*., 2021). Our time lapse imaging and the cell painting assay revealed additional evidences of morphological and biochemical hallmarks of ferroptosis (Dixon *et al*., 2012; Tang *et al*., 2021; Chen *et al*., 2021a, 2021b) and our pathway enrichment analysis from the screening assay provided additional evidence. A significant number of ferroptosis inhibitors were among our primary hits whereas apoptosis and autophagy inhibitors were absent. Finally, the two-fold enrichment in the percentage of ferroptosis inhibitors among our active hits between the primary screen and the cell painting assay consisted of a good validation for the cell death mechanism as well as for the screening pipeline.

Since its first description a decade ago, ferroptosis (Dixon *et al*., 2012) has gathered much attention as it is implicated into a diverse group of age-related diseases like cancer, cardiovascular diseases, stroke and NDD (Zhou, 2020). Increasing evidence in the literature links ferroptosis in the pathophysiology of tauopathies and other NDD (Zhang *et al*., 2018; Masaldan *et al*., 2019; Ndayisaba *et al*., 2019; Angelova *et al*., 2021). After all, iron accumulation and lipid peroxidation in postmortem brain tissue of patients with NDD are known long before the definition of ferroptosis (Angelova 2021). At the molecular level, iron co-localizes with Aβ plaques, promotes neurofibrillary tangles (NFT) through Tau hyperphosphorylation and induces nitration of Tau, which prevents microtubule stabilization (Masaldan *et al*., 2019; Ndayisaba *et al*., 2019; Reichert *et al*., 2020). In parallel, lipid peroxidation another biochemical hallmark of ferroptosis (Hong-fa Yan *et al*., 2021) and the first principal component feature defining our phenotype of MCOS in cell painting assay, is evident in numerous NDD, including tauopathies (Zhang *et al*., 2018; Bartolome *et al*., 2022). Lipid peroxidation products contribute to MAPT phosphorylation in primary cultures of mouse embryo cortical neurons (Gómez-Ramos *et al*., 2003), while are selectively associated with PSP via NFT formation (Odetti *et al*., 2000). Elevated levels of lipid peroxidation, ROS, and mitochondrial dysfunction were also detected in mesenchymal stem cells derived from patients with PSP compared to controls (Angelova *et al*., 2021).

Taken these together, we postulate that ferroptosis is induced under MCOS conditions in differentiating iNGN2 cortical neurons and their vulnerability depends on their genetic background of MAPT-17q21.31 locus haplotypes, with the risk allele H1/H1 being more sensitive. More detailed work needs to be done in order to address the causative root of OS induced ferroptosis in tauopathies and other NDD but always in the light of MAPT -17q21.31 locus haplotypes including the enrolment of other genes within the locus.

After elucidating the difference to AO depletion between the MAPT-17q21.31 locus haplotypes, we implemented a screening pipeline to identify candidate molecules that could rescue and/or reverse the lethal effect of MCOS on sensitive H1/H1 neurons. 6.7% of our initial FDA-library returned as primary hits with several of them being reported earlier as drugs that either prevented iron (Fe2+) induced neurotoxicity, or they were documented as ferroptosis inhibitors (Faissner *et al*., 2017; Stockwell *et al*., 2017; Mou *et al*., 2019; Tang *et al*. 2021). Since, AO defense systems is a complex, multi-molecular regulatory pathway with enzymatic and non-enzymatic AO, inhibitors, cofactors, scavengers, and metal chelators (Kyung *et al*., 2020; Lee *et al*., 2020) it was not surprising that we had a quite functionally and structurally diverse group of small molecules as primary hits. Structure similarity analysis returned twelve clusters of primary hits with polyphenolic molecules grouped in clusters 1 and 2, Vitamins E and K in cluster 12 and AO analogues in clusters 10 and 11. Small clusters contained functionally and structurally related drugs with similar therapeutic uses. For example bisphosphonates of Cluster 3, used in bones treatment (Allgrove, 1997; Vasikaran, 2001; Roelofs *et al*., 2010), are reported to retain AO properties by either inhibiting the lipid peroxidation and the Fenton reaction and/or due to their calcium chelating activity (Rogers *et al*., 2000; Villegas *et al*., 2014).

An initial inspection of molecular targets of primary hits, according to Selleckchem, demonstrated a wide range of molecular activities and a weak association of primary hits with the AO pathway. In our pathway enrichment analysis, we employed two complementary approaches, by relying on drugs-proteins interactions from DrugBank and interactions mined from the literature. The DrugBank interactions results were in agreement with the molecular targets provided by Selleckchem where the most enriched pathway was the *G alpha signaling events* due to agonist/antagonist of G-protein coupled receptors among our primary hits, but were unrelated to MCOS and the ferroptosis phenotype. In contrast, the text mining approach, that leverages a wider and more up to date range of interactions mined from the literature revealed as the most targeted pathways *lipid metabolism* and *biological oxidation*. One of the main reasons for this discrepancy is probably that most of the more recent drug-protein interactions from the literature are not yet annotated in databases. Additionally, as none of the three major databases which we used included a ferroptosis related pathway, we integrated the FerrDb in the analysis. Based on DrugBank interactions, primary hits were not associated to any kind of cellular death mechanism, as opposed to the text mining interactions where both the ferroptosis and the apoptosis pathways were targeted by more than the half of our primary hits, verifying further our phenotype and the screening assay results.

In the next steps of validation and confirmation of primary hits, we employed a secondary screen with DRC and TAL analysis and further excluded a number of primary hits from the list. In DRC analysis, we did not obtain any false positive hits, as all molecules were still active at 5 μM. The n=4 replication, the proper z-score cut off in conjunction with the z-score standard deviation inclusion in the selection criteria of primary hits, could be the explanation. Several single dose active molecules were excluded from further analysis and one of them was Minocycline, a second-generation tetracycline (TC) which showed high efficacy only at 5 μM in our primary and secondary screens. Our initial library included other TCs with some potency in reversing the neurotoxicity but all were below the cut off used in the primary screen. It is known that TC have direct scavenging activity towards free radicals (Kładna *et al*., 2011) and Minocycline was found to be neuroprotective in experimental ischemic stroke model and attenuated Iron-Induced Brain Injury (Matsukawa *et al*., 2009; Zhao *et al*., 2016). In order to detect possible primary hits which could have retained live neuronal body counts but had degenerated axons, we did a retrospective TAL analysis of the DRC experiment and further eliminated primary hits from the list. One of them was the β- blocker Carvedilol, which has been shown to be the most potent scavenger of free radicals among other β-adrenoceptor blockers. The IC50 of Carvedilol in inhibiting (Fe2+)-initiated lipid peroxidation in rat brain homogenate was determined as 8.1 μM. (Yue *et al*., 1992), 3.1 μM higher than the concentration that we used. This could be the possible explanation why Carvedilol was not efficient enough to rescue the MCOS induced ferroptosis in iNeurons by retaining intact the axonal network and also could not reach out the highest asymptote at 5 μM concentration in DRC experiments.

Lastly, OS caused damage to nucleic acids, proteins and lipids is known for a very long time (Lassmann and van Horssen, 2015). In order to validate the health status of neurons at subcellular level after being rescued by the active hits we used Cell Painting assay and imaged live neurons under MCOS during the dying process in real time. PCA analysis of features extracted from Cell Painting assay helped us not only to exclude further nine drugs from the active hits list but also confirmed the ferroptosis as the cellular death model induced by our MCOS phenotype. All the characteristic features of ferroptosis, smaller-destroyed mitochondria, plasma membrane rupture and round up of cells (Dixon *et al*., 2012; Tang *et al*., 2020; Chen *et al*., 2021a, 2021b) were detected during the assay. The lipid peroxidation level, one of the main indicators of ferroptosis (Hong-fa Yan *et al*., 2021), based on the PCA was the most significant feature that defined PC score 1 on day 8 neurons. While the sizes and the numbers of lysosomes defined the PC score 3 on day 9 neurons is in good agreement with the results of Tian *et al*., (2021) who demonstrated a strong correlation between lysosomal failure and ferroptosis in a genome wide CRISPRa/i study in human neurons under OS due to AO depletion.

MAPT, the most studied gene of the MAPT-17q21.31 locus with alternatively spliced isoforms, it is known for its significance in neuronal development, functioning, microtubule organization and axonal trafficking as well as for its contribution to numerous NDD including tauopathies (Goedert *et al*., 1989; Andreadis *et al*., 1992; Liu and Gong, 2008; Chung *et al*., 2016; Park *et al*., 2016; Prezel *et al*., 2017). Since evidence of axonal blebbing/swelling, microtubule bundle dis-arrangement and axonal degeneration preceded the neuronal death induced by MCOS, we took a closer look to the MAPT region within the locus. We hypothesize that possible explanations for the difference in resistance between the haplotypes could be the haplotype defining SNPs reported previously as modulators of MAPT isoforms expression in two independent studies (Lai *et al*., 2017; Wang *et al*., 2016). SNPs, rs17651213 and rs1800547 were proven to regulate the haplotype-specific splicing of MAPT isoforms under basal conditions by whole-locus genomic DNA expression vectors (Lai *et al*., 2017). While SNPs rs241032 and rs242561 were determined in an association study of genome-wide mapping of NRF2/sMAF occupancy with disease risk SNPs identified in GWAS (Wang *et al*., 2016). The former SNP rs241032 (downstream of CRHR1-IT1) was not found among the NRF2/sMAF-occupied genomic regions based on previous ChIP-seq data, in contrast with the SNP rs242561 which is located on a cis-acting enhancer of anti-oxidant response element (ARE) on intron 1 of MAPT. According to the study the C/T conversion in H2/H2 haplotype confers stronger binding of the major AO transcription factor Nrf2 which is already proven to have essential role in ferroptosis (Anandhan *et al*., 2020; Sun *et al*., 2020; Zhou *et al*., 2020; Chen *et al*., 2021). Stronger affinity of Nrf2 to the ARE element in H2/H2 alters MAPT expression in neural cells and in cerebellum of Nrf2 mouse OS model. Although the study was not conclusive whether this in vivo regulation of MAPT was a basal effect or induced by OS conditions, additionally none of their models included the H2/H2 haplotype and the stronger T allele (Wang *et al*., 2016). Further in detailed studies needs to be done in order to validate at molecular level the contribution of MAPT together with the other genes of the MAPT-17q21.31 locus to the differential resistance of haplotypes to MCOS induced ferroptosis.

## Conclusion

The MAPT H1/H1 haplotype but not the H2/H2, consists of a common risk factor for various NDD including important tauopathies. Here, we presented experimental evidence that the H1/H1 haplotype exhibits an increased sensitivity to ferroptosis induced by MCOS compared to the H2/H2 haplotype. Ferroptosis induced by AO depletion was observed only at neuronal maturation stages with complex axonal networks without affecting the smNPC’s and the initial stages of neuronal differentiation. It caused microtubule bundles disarrangements and axonal degeneration prior to neuronal death in the same manner on both haplotypes, but earlier on H1/H1 haplotype. We identified FDA-approved drugs that possess AO activity and that were effective in inhibiting ferroptosis in a dose dependent manner.

Despite the huge number of AO enrolled in clinical trials against NDD, factors like the bioavailability, instability, limited transport and targeting the right ROS species have been attributed so far to their controversial outcomes. Based on our results, additional reasons could be that the recruitment and the results of these studies are mostly evaluated based on clinical symptoms while the genetic background and the genetic risk factors are being overlooked. Additionally, we showed that AO supplementation could inhibit ferroptosis at the stage of axonal degeneration which precedes neuronal death. In humans these two events occur before the onset of clinical symptoms whereas, the eligibility in the majority of the clinical trials is based on clinical symptomatology and already diagnosed patients with NDD. Finally, although OS causes simultaneous damage to several types of macromolecules, most of the clinical trials are conducted with one AO molecule which is targeting only a specific outcome of OS.

Future studies need to shed light and pinpoint the possible SNPs responsible for the difference in between the MAPT haplotypes susceptibility to MCOS, while uncovering in detail the steps of ROS - ferroptosis induction on neurons from patient-derived cell lines.

## Materials and Methods

### Cell Culture – Mild Chronic oxidative stress model on iNeurons

smNPC derivation from iPS and expansion was done according to P. Reinhardt *et al*., 2013. smNPC were transduced with Neurogenin 2 (NGN2) containing Lentivirus (Dhingra A. *et al*., 2020) and differentiated onto Poly-L-Ornithine / laminin coated 96 black well plates (PerkinEnmayer) with 2.5 μg/mL doxycycline and 10 μM DAPT according to others (Topol. A *et al*., 2015). Complete media changes were performed on days 3 and 6 of neuronal differentiation with N2B27 media supplemented with Dox, DAPT and 10 ng/mL brain-derived neurotrophic factor (BDNF), 10 ng/mL glial cell-derived neurotrophic factor (GDNF), 10 ng/mL neurotrophic factor 3 (NT-3), 0.5 μg/mL Laminin. Only half media changes were done for longer neuronal differentiation. Neuronal media changes were done with VIAFLO electronic pipette (INTEGRA) with the minimum flow rates on individual plates. All cell lines were fully characterized in previous study (Strauß T. *et al*., 2021).

MCOS conditions were induced by culturing smNPCs and differentiating into neurons with B27 supplement minus Antioxidant (10889038, Gibco). MCOS rescue experiments were done on day 6 of neuronal differentiation by replacing the minus AO with plus AO media or drugs treatment.

In all experiments the outer wells of 96 well plates were not used to avoid the edge effect. Viability assays were done with CellTrace™ Calcein Green or Red-Orange (C34852, C34851, ThermoFisher) and Spectramax M2 Multimode Microplate Reader (Molecular Devices) or Cell Voyager Yokogawa (CV700). For the time lapse images of ferroptosis propagation LIVE/DEAD™ Viability/Cytotoxicity Kit (L3224, ThermoFisher) was used.

### Screening Assay set up

In order to establish the screening assay conditions, we had to specify different parameters. For the screening assay we used the smNPC, H1/H1 diku line treated without AO for five weeks, differentiated into neurons in minus AO media and the rescue conditions and imaging time points were established (**Fig.S2B** and **S2C**). Results are the mean ± SEM from three replicates. In comparison with One-Way ANOVA of groups with d3 and d6 neurons Initially, we determined day 6 as the latest time point in reversing the neurotoxicity by AO supplementation, since there was no significant difference in survival of day 3 and day 6 rescued neurons at all four imaging time points (**Fig.S2B**). We also tested different imaging time points (day 8, day 10, day 12 and day 14) and the best was determined to be day 12 of differentiation, where the difference in numbers of live to dead neurons between day 12 rescued versus not rescued was significantly the highest (**Fig.S2B**). Additionally, we observed that the rescue with AO supplementation upon day 6 of differentiation was dose dependent and even low concentrations (1/625 dilution) of AO could rescue the phenotype moderately (**Fig.S2B**).

Our library of small molecules was aliquoted and stored at 1mM in DMSO. Previous studies have shown the cytotoxic effect of increasing doses of DMSO on primary neurons (Zhang *et al*., 2017). In order to determine the highest tolerable concentration by our stressed neurons, we rescued the neurotoxicity with plus AO on day 6, including or not 0.5% and 1% of DMSO. Based on results (**Fig.S2C**) some additional toxicity with only 1% DMSO and not with 0.5% was observed on days 10, 12 and 14 of differentiation. Finally, we evaluated the performance of our screening conditions with the Z-factors (Zhang *et al*., 1999; Brideau *et al*., 2003) for each imaging time point. The best Z-factors of 0.89 and 0.88 were calculated for imaging days 12 and 14 respectively.

### Primary and secondary screen of small molecules

The FDA-approved Drug Library (L1300, Selleckchem) was obtained by Hoelzel diagnostika and diluted to 1 mM in DMSO, aliquoted in 96 deep well plates and stored at −80 °C. Compounds plates were thawed and equilibrated to room temperature before use. Screening plates were handled in a semi-automated system with VIAFLO 96/384 handheld electronic pipette (INTEGRA) and Multi-Mode Dispenser MultiFlo FX (BioTek). Screening assay was performed with H1/H1 line diku treated and differentiated into neurons with standard day 3 media change with minus AO media. Upon day 6 of media change, minus AO was replaced with plus AO media as positive control whereas minus AO media was used as negative control. Both controls include 0.05% DMSO. Primary screening was performed in two independent batches and in total 1430 compounds were screened in n=4 replicates. The highest concentration of small molecules was 5 μM since neurons were sensitive to higher concentrations of DMSO as determined by screening assay set up. At the end of the treatment live neuronal staining was performed with Calcein Red-Orange addition to the media and further incubation for 30min at 37°C. Each batch included 60 plates with 3,600 wells and 15 fields per well were imaged on Cell Voyager 7000 (Yokogawa). Live neuronal counts were analyzed with open-access software for High Throughput Screening application HitSeekR (List *et al*., 2016) using plate-wise normalization.The replicates correlation with corresponding R^2^ from the analysis are shown also (**Fig.S2E**)

Dose response curve experiment was done by cherry picking manually of primary hits from the library and reformatting into two deep well plates with minus and plus AO controls included. Five-fold serial dilutions (from 5μM to 8nM) were prepared into new deep well plates and transferred to neurons during day 6 media change with VIAFLO 96/384. Cell staining and imaging was performed as before.

### Pathway enrichment analysis

We employed two complementary approaches to identify chemical-protein interactions of primary hits. In the first approach, we relied on well-established interactions from DrugBank that were further enriched via manual curation. On the other hand, the second approach employed a text mining engine (SCAIView; https://academia.scaiview.com) to systematically extract chemical-protein interactions described in scientific literature. The complementarity of both approaches enabled us to not only rely on well-studied interactions, but also leverage other interactions mined from the literature (**Fig.3 step 1**). Next, by using the interactions retrieved from both approaches we mapped proteins that interact with each of the small molecules to pathways in three major pathway databases (i.e., KEGG, Reactome, and WikiPathways) (**Fig.3 step 2**). Since these databases contain multiple equivalent pathways, they were grouped together using their pathway hierarchy as well as the mappings from (Domingo-Fernández *et al*., 2019). Finally, we compared the most common pathways targeted by our list of primary hits from each of the two approaches (i.e, text mining and DrugBank) in order to identify the set of pathways targeted by these small molecules (**Fig.3A step 3**). Additionally, we included the pathway representing the newly described cell death mechanism ferroptosis using the gene set curated from FerrDb as it was not integrated into any of the three major pathway databases.

In parallel, we checked the number of our primary hits that have already been investigated in the context of neurodegenerative disorders, querying ClinicalTrials.gov using all names and identifiers of the 94 diseases belonging to the MeSH branch of “Neurodegenerative Diseases” (date of the query 13.05.2021). The resulting query revealed that 34 small molecules of our primary hits have already been investigated in a clinical trial for any neurodegenerative disease (data not shown).

### Image Acquisition and Analysis

Fluorescence image acquisition throughout the study was done on Cell Voyager spinning disk confocal CSU imaging system CV7000 (Yokogawa) equipped with a stage incubator which provides long-term time lapse imaging of live cells. Stage incubator with 5% CO_2_ and 37^0^C temperature was used for all imaging experiments.

Primary screen was imaged with 10× air objective in single z-stack and 15 fields per well, producing in total 900 images per plate in seven minutes and 54.000 images per run. In secondary screening of dose response curve experiment 14 fields of single z-stack were used per well for all concentrations except the highest of 5 μM which was imaged as the maximum projection of two z-stacks in order to be used in TAL analysis.

Time Lapse images of 5 and 24 hours were imaged every 20 and 60 minutes respectively.

Throughout the study image analysis was done with CellPathfinder v 3.03.02.02 and the blebs counting in Cell Painting assay was performed with v 3.06.01 which includes deep learning functions. Only for the neuronal counts of dose response curve experiments the CellProfiler (version 2.2.0) was used due to lack of nucleus staining and extensive clumping of cells. Whereas ImageJ (2.3.0/1.53f51, Java 1.8.0_172 (64-bit)) was used occasionally to construct bright field image composites.

### Cell Painting Assay

After 48h and 72h of incubation with the compounds, cells were stained with the fluorescence probes indicated in Table 1 upon day8 and day9 of differentiation. The staining solutions were prepared as 4X concentrations in minus AO media, added to each well (50 μL) and incubated at 37°C for 30 min. After incubation and before imaging half of the media was replaced. Each time point was carried out as n=4 of replicate plates. The combination of staining’s per day were determined in a preliminary test based on our MCOS induced ferroptosis phenotype, the cell type lineage and the experimental parameters as fixing, staining and subsequent imaging of stressed neurons was not possible. Previously, we observed that the very first cellular compartments which were affected and destroyed were the mitochondria and the cellular membrane lipids. We set up the imaging of the above cellular compartments alongside nucleus and microtubules as reference, on day8 of differentiation, which is the earliest time point we observe stressed neurons. For day 9 neurons where we observed more advanced stages of axonal degeneration with extensive blebbing and dying neurons in minus AO controls we used fluorescence probes for cell membrane integrity, lysosomes, nucleus and microtubules.

**Table 1.**
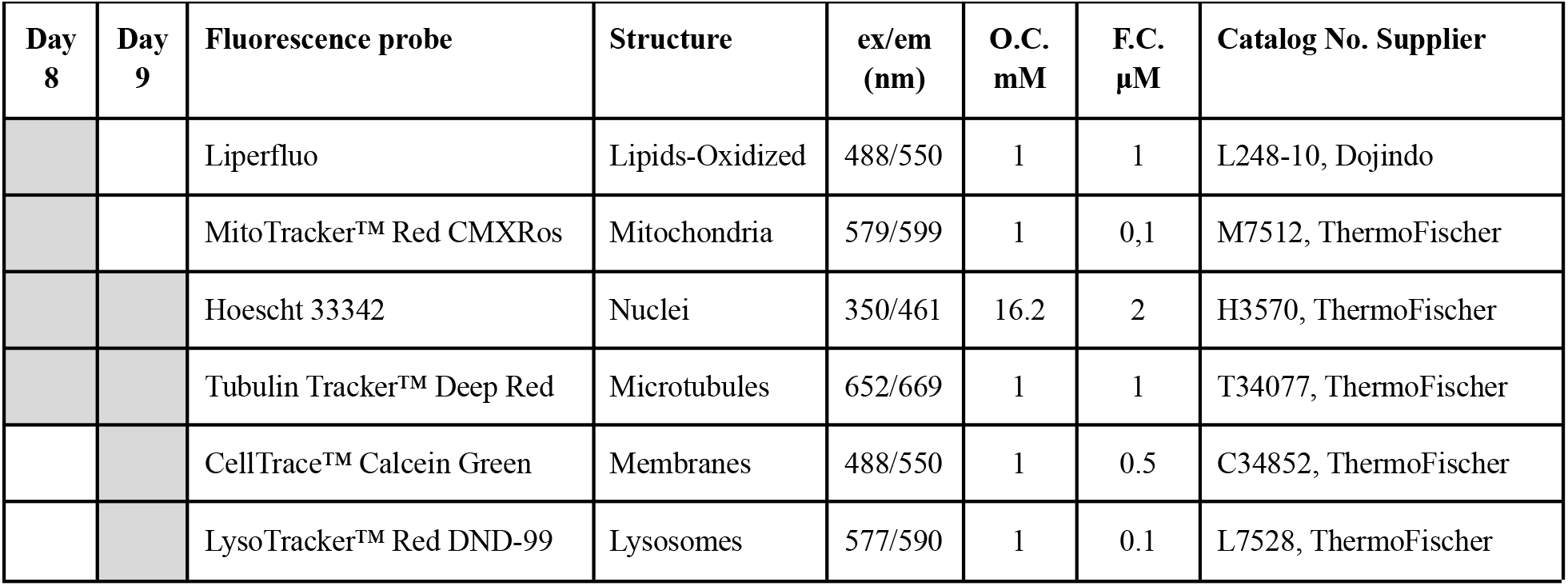
Cell Painting Fluorescence probes.

Imaging conditions were set up with the aim to cover the maximum number of fields per well in less than 2h per plate, as we were imaging on real time degenerating neurons during the dying process which did not allow us to perform fixation. Cell Voyager technology equipped with a stage incubator for live cell imaging, we imaged 14 fields per well with 60X water immersion objectives. Big objects like nucleus, cell body and cell membranes were imaged as Sum of z-stacks projections whereas small objects like mitochondria and lysosomes were imaged as the Maximum of z-stacks projections.

Dark and shading correction of images were done with CV Image Correction Tool R2.02.06. Cell Pathfinder (CPF) software was used for subsequent image analysis. Initially, blur or damaged images were removed from further analysis by using the Thumbnail function of CPF and simultaneously examination in a plate-wise manner of all fields. Machine learning function was used to identify all nucleus from the Hoechst channel and by applying filters they were separated to Live and Dead Nucleus (NucL. and NucD.) categories. The live nucleus seeds were used as a reference region to identify the cell boundaries for the lipids and the oxidized lipids staining on day 8 and the cell membranes on day 9. Mitochondria of neuronal soma were excluded from detection and only mitochondria on axonal areas were determined as the cytoplasmic mitochondria of neurons are difficult to segment and need different imaging strategies. Lysosomes, on day 9 neurons, were identified and separated into axonal and cell body lysosome groups. For both days of imaging, a deep red channel was used to detect the axons/microtubules. The axonal length and the area covered by axon related features were used in the analysis, as the junctions and branching counts cannot be determined accurately in cultures with advanced neural networks. Finally, day 9 images were used for the identification and the counting of blebs on axons with deep machine-learning function and deep image gating with CellPathfinder (ver. 3.06.01).

The above determined objects were used to measure and extract morphological features related to size, shape, texture and intensity per field and per object. Additional mathematical expressions like the percentage of live nucleus counts versus the total number of nucleus identified were extracted and used for the analysis and the quality control in comparison between and among the plates. Pearson correlation was determined with positive controls among the plates to exclude possible batch effects as part of quality control as is suggested in Caicedo *et al*., 2017. In general, we extracted and used in further dimensionality reduction analysis with Principal Component Analysis (PCA) in total 200 features for day 8 and 206 features for day 9 after removing all the total values which were not averaged either per field or per cell number.

### Research Tools and Software

Statistical analysis was done with Excel (Microsoft Corporation. (2018). Retrieved from https://office.microsoft.com/excel) and GraphPad Prism version 9.1.0 for Windows (GraphPad Software, La Jolla California USA, www.graphpad.com). All figure composites were done with Biorender (Created with BioRender.com). Primary screening analysis was done with the open-access software for High Throughput screening application HitSeekR (List *et al*., 2016). The chemical structure similarity was performed with the Java-based open source tool for the visual analysis and interactive exploration of chemical space Scaffold Hunter 2.6.3 (Schäfer *et al*., 2017). Cell Painting assay analysis and plots were created by using R (http://www.R-project.org) and packages FactoMineR (Sebastien *et al*., 2008), Factoextra (Kassambara and Mundt 2020), ggplot2 (Wickham, 2016), rgl (Murdoch and Adler, 2021) and magic (Ooms, 2021).

## ACKNOWLEDGEMENTS

We would like to acknowledge the assistance of Dr. Joachim Täger for the plate reformatting of the initial FDA-approved chemical library and Mahomi Suzuki, Senior Application Specialist in Global Sales Department of Life Business HQ, Yokogawa Electric Corporation 2-3 Hokuyodai, Kanazawa-shi, Ishikawa, 920-0177, Japan for the detection of axonal blebs with deep learning function in Cell Painting assay.

## Author Contributions

E. Sadikoglou: conceptualization, data curation, analysis, investigation, methodology, writing original draft, review and editing. Domingo-Fernández D, Kodamullil A: Bioinformatic analysis

A. Illarionova: PubChem small molecules compound ID’s

T. Strauß: iPSC to NGN2_smNPC derivation

SC. Schwarz, GU. Höglinger: conceptualization,, supervision, review

A. Dhingra: primary screening methodology, review and editing

P. Heutink: conceptualization, resources, supervision, review and editing.

All authors commented and approved the final version of the manuscript.

---

**Fig. S1.**
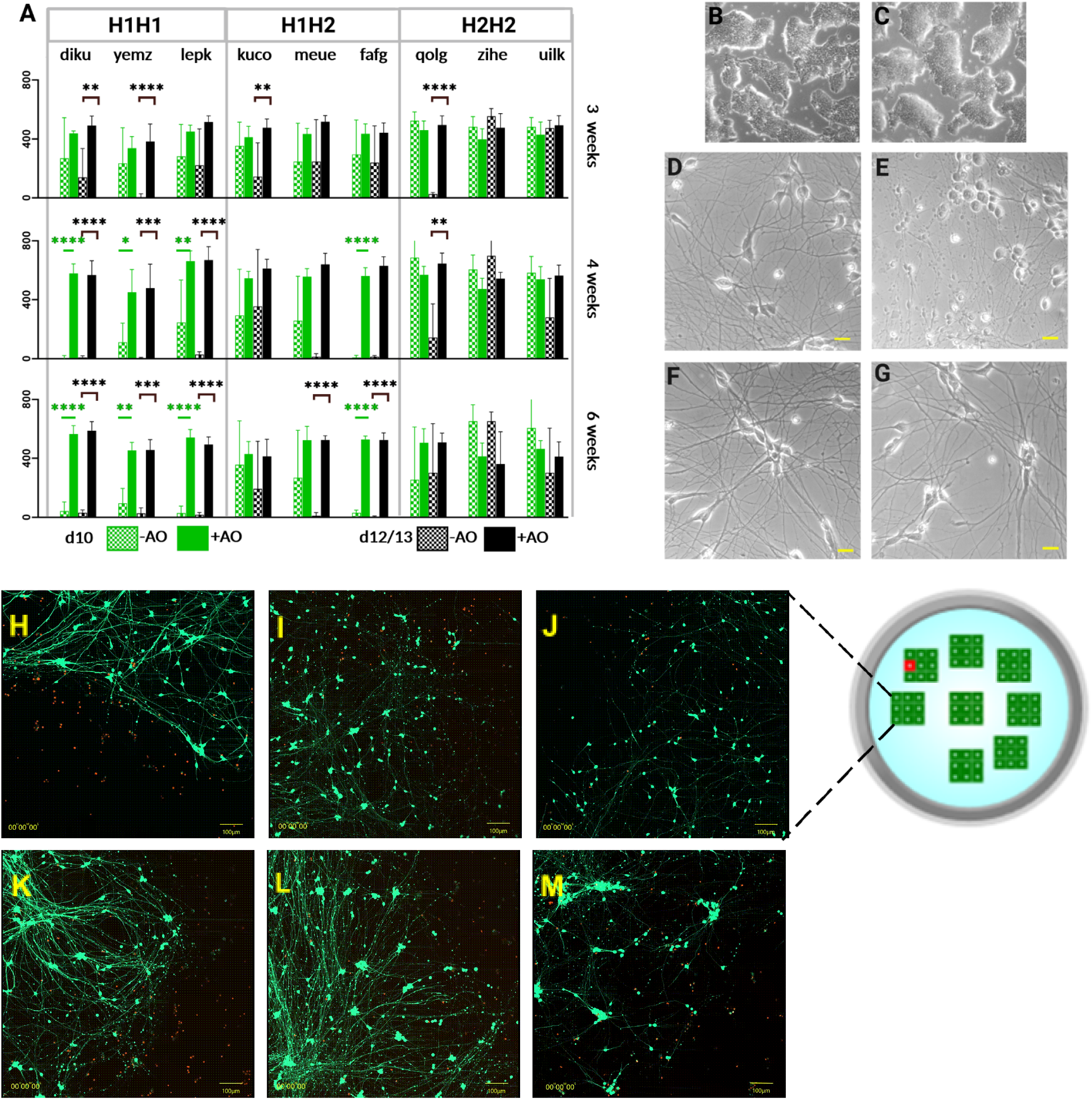
Diffèrence in resistance of MAPT haplotypes neurons to ferroptosis due to MCOS. **(A)** Three smNPCs per haplotype were treated with and without AO for 3,4 and 6 weeks. Relative fluorescence units (RFU) of iNGN2 neurons stained with Calcein -Red were determined. Data represent mean ± SD of normalized RFU-Statistical analysis was done with two -tailed Student’s t tests and P< 0.05: *, P< 0.01: **, P< 0.001: *** P< 0.0001 ****. **(B-C)** Representative bright field images of smNPC Hl/Hl (diku) treated for 5 weeks with and without AO. **(D-E)** Hl/Hl (diku) and **(F-G)** H2/H2 (uilk) day 8 neurons from smNPC’s treated for 5 weeks with (left) and without AO (right). Axonal degeneration, blebs formation and round up of neuronal soma observed in Hl/HI diku **E**. Scale bar 100 pixels on ImageJ, **(H-M)** Propagating cell death in population. Time lapse for 24 hours imaged every hour of 3×3 tiled images and DEAD/LIVE kit (red/green). Upper panel (H-l-J) H1/H1 cell lines (diku, yemz, lepk) and lower panel (K-L-M) H2/H2 lines (qolg, zihe, uilk) treated for 5 weeks in minus AO media. H-I imaged on day9, J-K-L day 12 and M day 21 of neuronal differentiation. Scale bar 100μm. For complete time-lapse videos refer to **Videos S9-14** respectively.

**Fig. S2.**
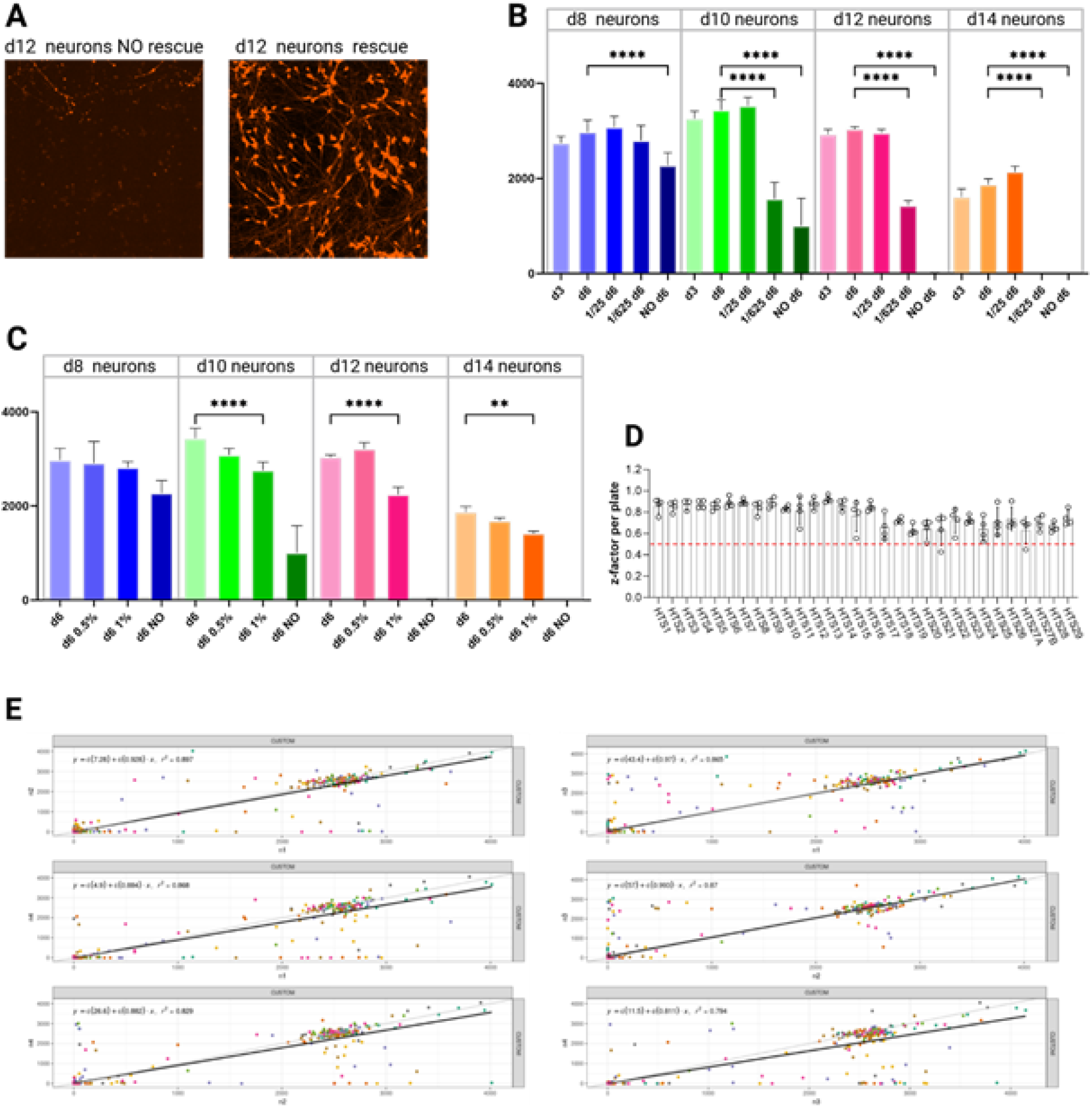
Primary screen conditions and quality control with H1/H1 line diku. **(A)** Representative images of day 12 neurons imaged with Calcein Red of negative (no rescue on day 6) and positive (day 6 rescue) controls. **(B)** Live neuronal count on days 8, 10, 12 and 14 of neuronal differentiation from diku smNPC’s treated for 5 weeks without AO and rescued on days 3 and 6 with plus AO. Data represent mean ± SEM from three independent experiments compared with One-Way ANOVAand P< 0.05:* P< 0.01 : **, P< 0.001 : ***, P< 0.0001: ****. On day 6 of rescue additionally the phis AO media was diluted 25 and 625 times with minus AO media and the dose dependent rescue effect Is obvious. **(C)** The maximum DM SO concentration which is tolerable by stressed neurons on day 6 of rescue was determined as 0.5% since the 1% of DMSO is significantly different from no DMSO added on day 6 rescue. **(D)** z-factors from n=4 replicates per small molecule plate calculated from positive and negative wells in the primary screen. **(E)** Replicate correlation plot including a linear regression (black line) with corresponding R2 correlation factor from primary screen analysis with HitSeekR (List M,, et al,, 2016).

**Table S1.**
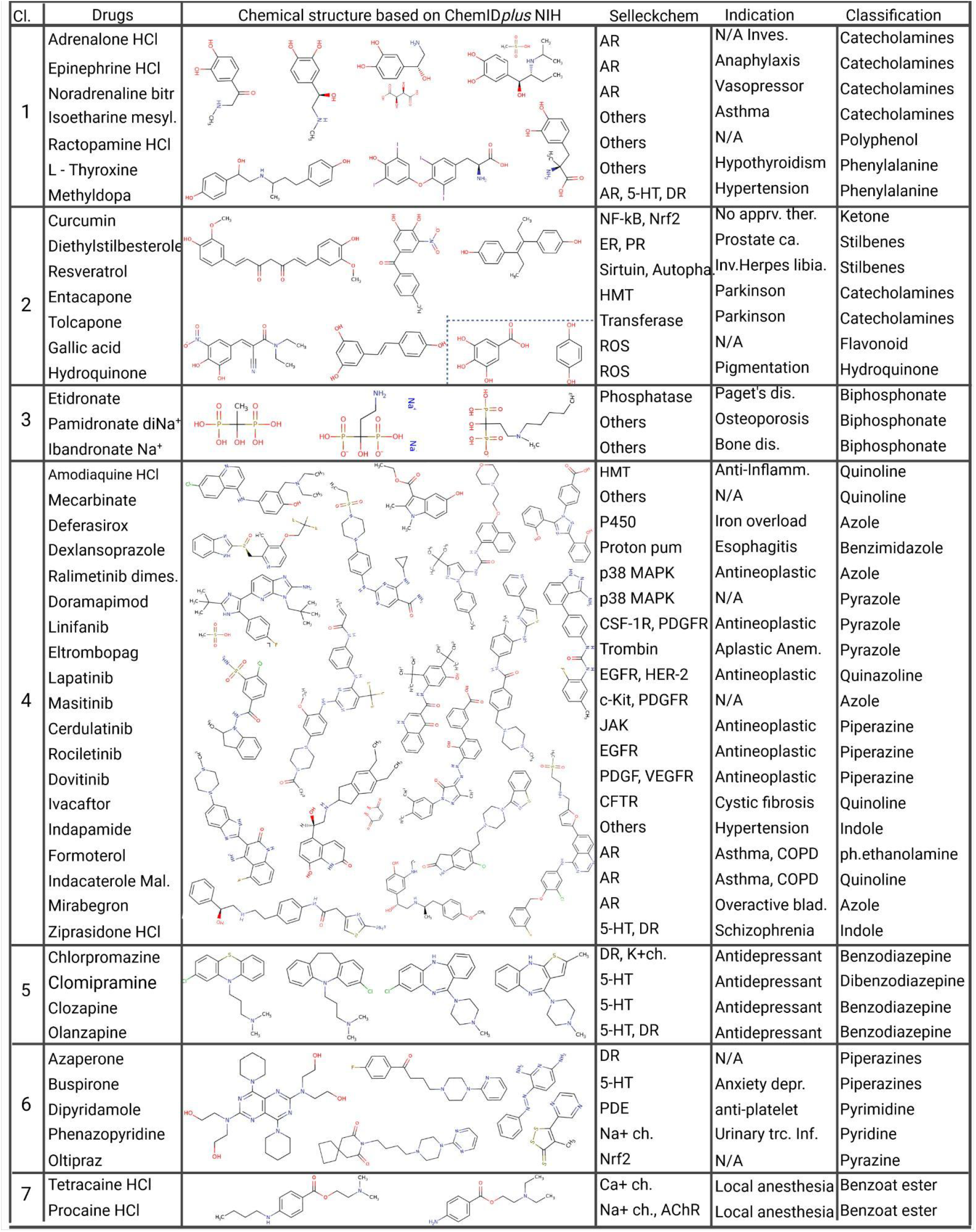

| Table S1 |  |  |  |  |  |
| --- | --- | --- | --- | --- | --- |
| Cl. | Drugs | Chemical structure based on ChemIDplus NIH | Selleckchem | Indication | Classification |
| 1 | Adrenalone HCl |  | AR | N/A Inves. | Catecholamines |
|  | Epinephrine HCl |  | AR | Anaphylaxis | Catecholamines |
|  | Noradrenaline bitartrate |  | AR | Vasopressor | Catecholamines |
|  | Isoetharine mesylate |  | Others | Asthma | Catecholamines |
|  | Ractopamine HCl |  | Others | N/A | Polyphenol |
|  | L - Thyroxine |  | Others | Hypothyroidism | Phenylalanine |
|  | Methyl dopa |  | AR, 5-HT, DR | Hypertension | Phenylalanine |
| 2 | Curcumin |  | NF-kB, Nrf2 | No appr. ther. | Ketone |
|  | Diethylstilbestrol |  | ER, PR | Prostate ca. | Stilbenes |
|  | Resveratrol |  | Sirtuin, Autophagy | Inv. Herpes libia. | Stilbenes |
|  | Entacapone |  | HMT | Parkinson | Catecholamines |
|  | Tolcapone |  | Transferase | Parkinson | Catecholamines |
|  | Gallic acid |  | ROS | N/A | Flavonoid |
|  | Hydroquinone |  | ROS | Pigmentation | Hydroquinone |
| 3 | Etidronate |  | Phosphatase | Paget's dis. | Biphosphonate |
|  | Pamidronate diNa<sup>+</sup> |  | Others | Osteoporosis | Biphosphonate |
|  | Ibandronate Na<sup>+</sup> |  | Others | Bone dis. | Biphosphonate |
| 4 | Amodiaquine HCl |  | HMT | Anti-Inflamm. | Quinoline |
|  | Mecarbinat |  | Others | N/A | Quinoline |
|  | Deferasirox |  | P450 | Iron overload | Azole |
|  | Dexlansoprazole |  | Proton pump | Esophagitis | Benzimidazole |
|  | Ralimetinib dimes. |  | p38 MAPK | Antineoplastic | Azole |
|  | Doramapimod |  | p38 MAPK | N/A | Pyrazole |
|  | Linifanib |  | CSF-1R, PDGFR | Antineoplastic | Pyrazole |
|  | Eltrombopag |  | Trombin | Aplastic Anem. | Pyrazole |
|  | Lapatinib |  | EGFR, HER-2 | Antineoplastic | Quinazoline |
|  | Masitinib |  | c-Kit, PDGFR | N/A | Azole |
|  | Cerdulatinib |  | JAK | Antineoplastic | Piperazine |
|  | Rociletinib |  | EGFR | Antineoplastic | Piperazine |
|  | Dovitinib |  | PDGF, VEGFR | Antineoplastic | Piperazine |
|  | Ivacaftor |  | CFTR | Cystic fibrosis | Quinoline |
|  | Indapamide |  | Others | Hypertension | Indole |
|  | Formoterol |  | AR | Asthma, COPD | ph.ethanolamine |
|  | Indacaterol Mal. |  | AR | Asthma, COPD | Quinoline |
|  | Mirabegron |  | AR | Overactive blad. | Azole |
|  | Ziprasidone HCl |  | 5-HT, DR | Schizophrenia | Indole |
| 5 | Chlorpromazine |  | DR, K+ch. | Antidepressant | Benzodiazepine |
|  | Clomipramine |  | 5-HT | Antidepressant | Dibenzodiazepine |
|  | Clozapine |  | 5-HT | Antidepressant | Benzodiazepine |
|  | Olanzapine |  | 5-HT, DR | Antidepressant | Benzodiazepine |
| 6 | Azaperone |  | DR | N/A | Piperazines |
|  | Buspirone |  | 5-HT | Anxiety depr. | Piperazines |
|  | Dipyrindamole |  | PDE | anti-platelet | Pyrimidine |
|  | Phenazopyridine |  | Na+ ch. | Urinary trc. Inf. | Pyridine |
|  | Oltipraz |  | Nrf2 | N/A | Pyrazine |
| 7 | Tetracaine HCl |  | Ca+ ch. | Local anesthesia | Benzoat ester |
|  | Procaine HCl |  | Na+ ch., AChR | Local anesthesia | Benzoat ester |

**Table S2.**
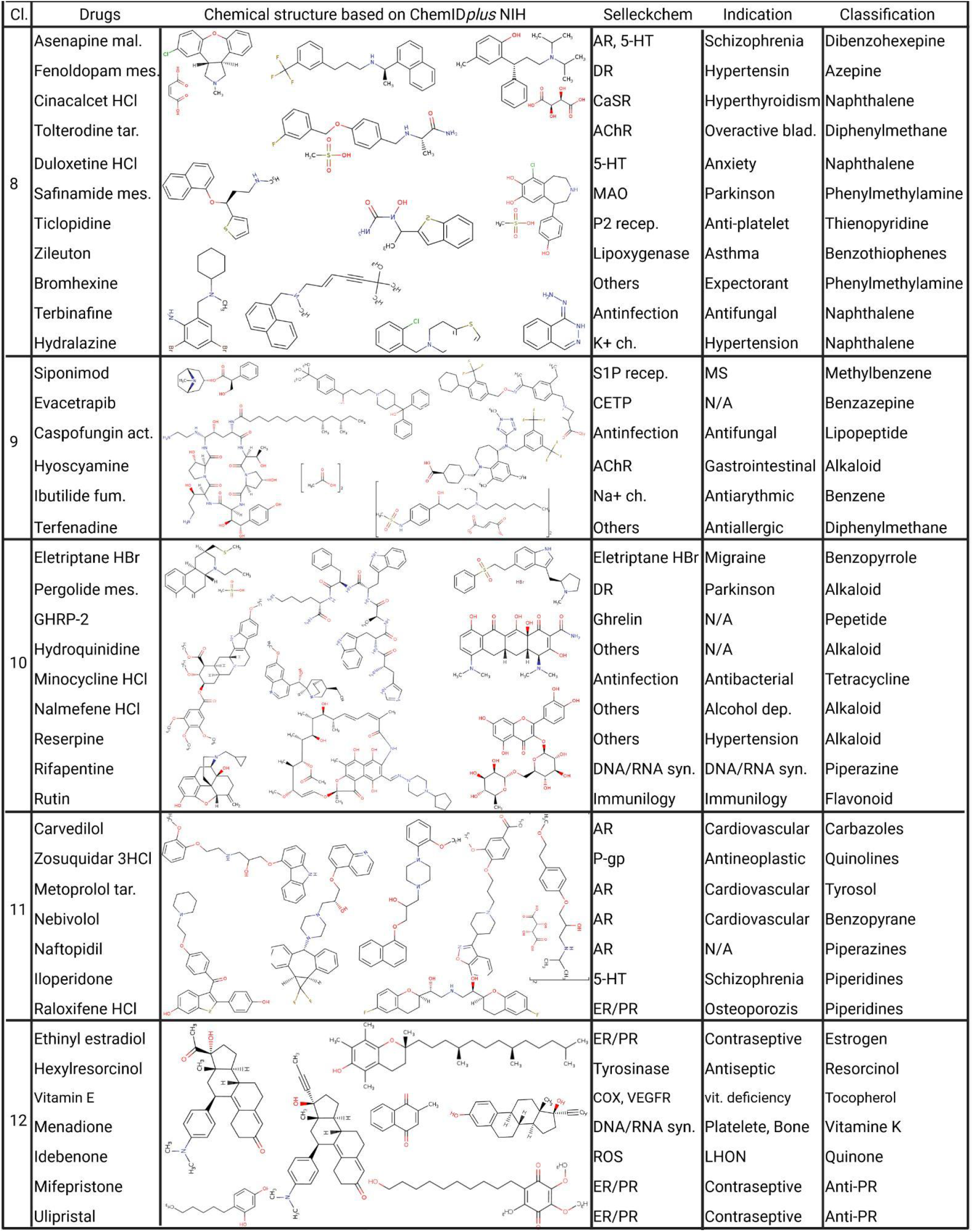

**Fig. S3:**
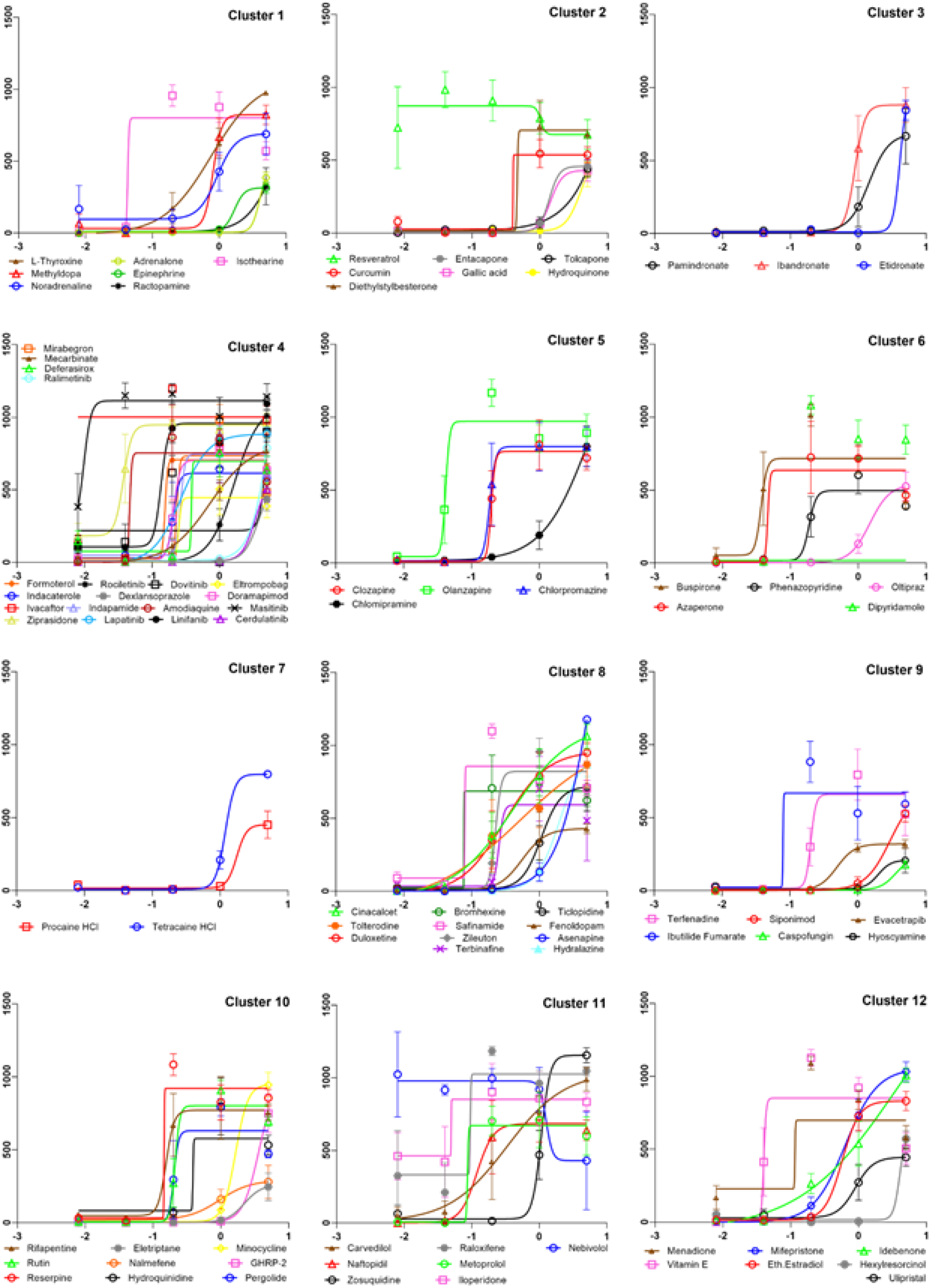
Dose Response Curves: Hits of primary screen were checked in a series of half-log concentration intervals, starting with the highest of 5μM and the lowest 8nM. Drugs were added as before on day 6 of neuronal differentiation on diku treated for five weeks without AO. Results were plotted per chemical structure similarity clusters in log scale of concentration indicated on x axes and live cell counts on y axes. Data points are the mean ± SEM from n=4 replicates.

