## Supplementary figures and images for "Rescue of the increased susceptibility to Mild Chronic Oxidative Stress of iNeurons carrying the MAPT Chromosome 17q21.3 H1/H1 risk allele by FDA-approved compounds"

### S1 video

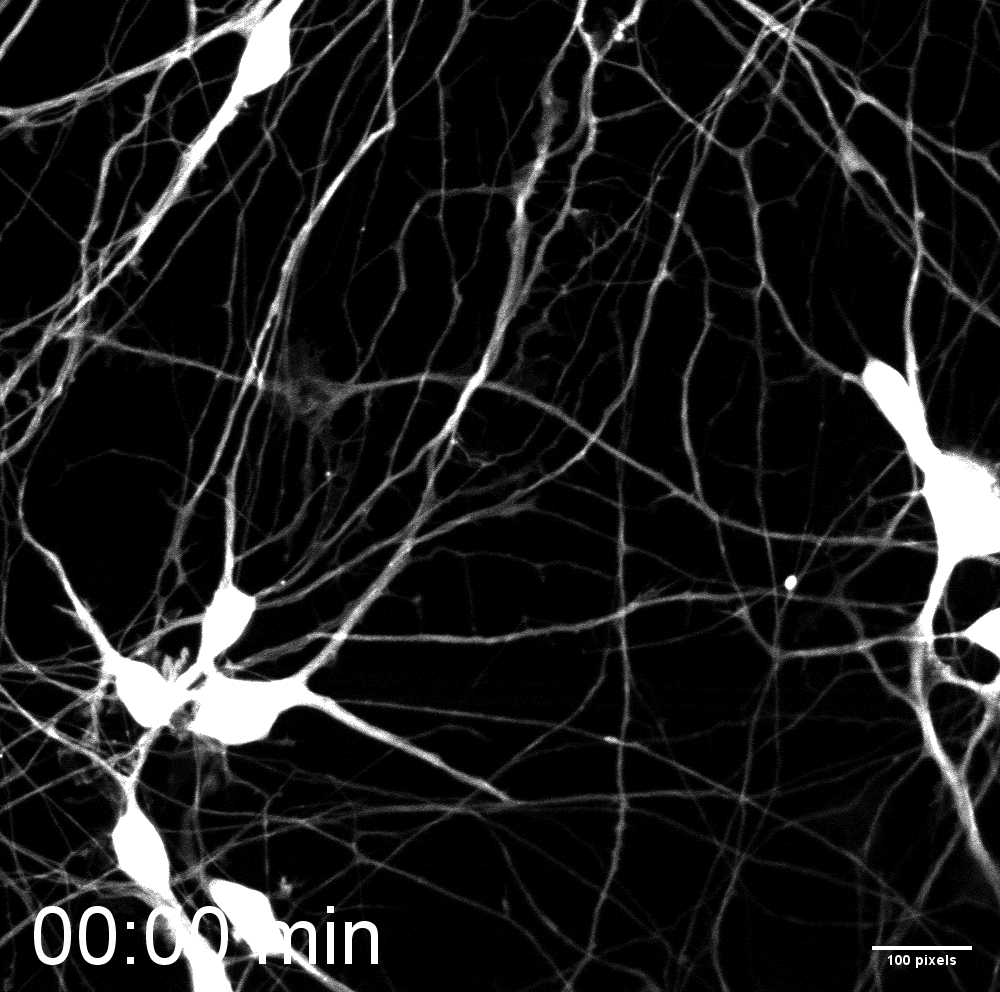

### S2 video

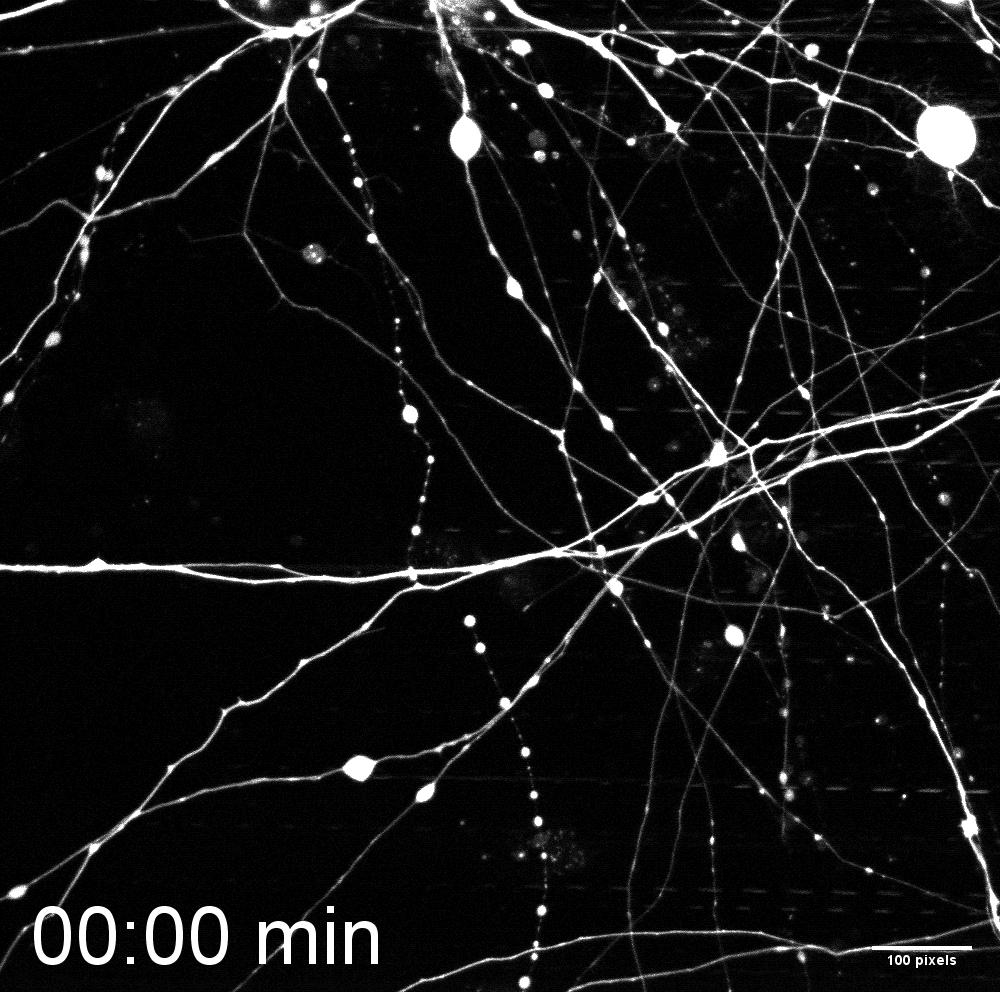

### S3 video

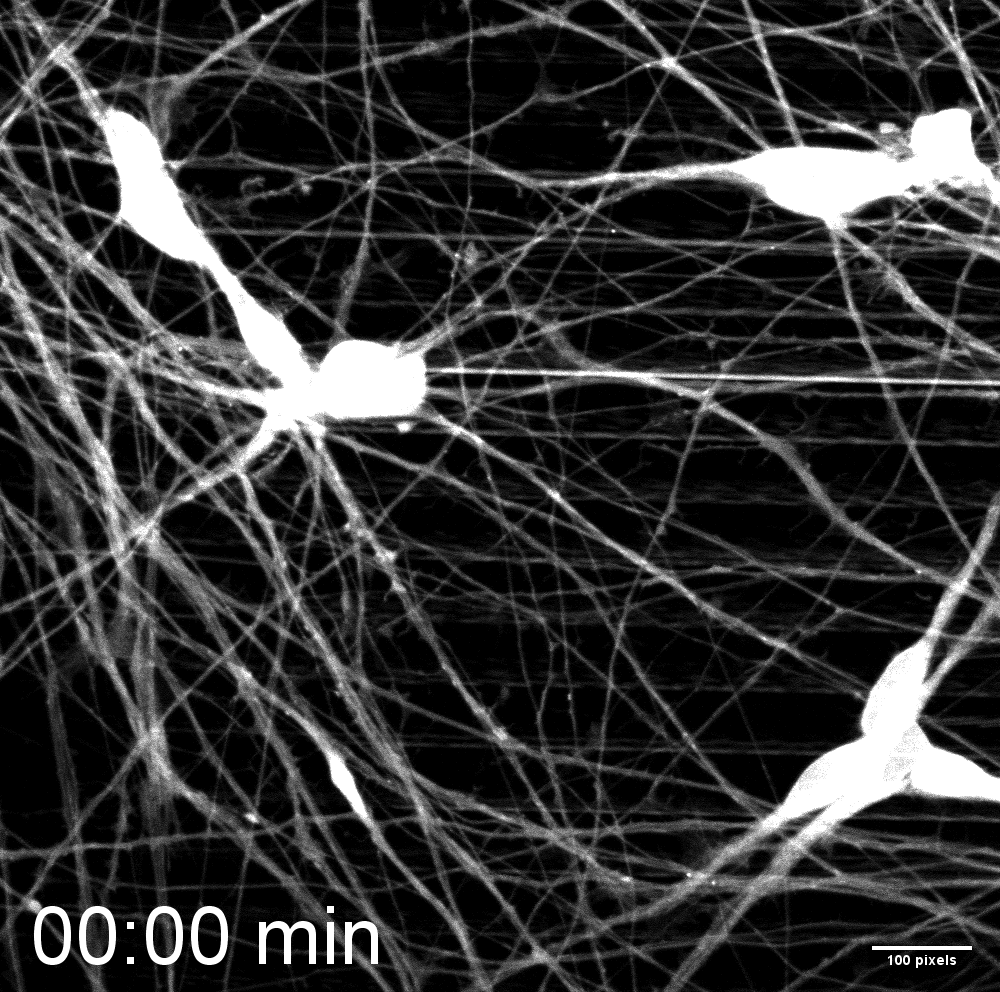

### S4 video

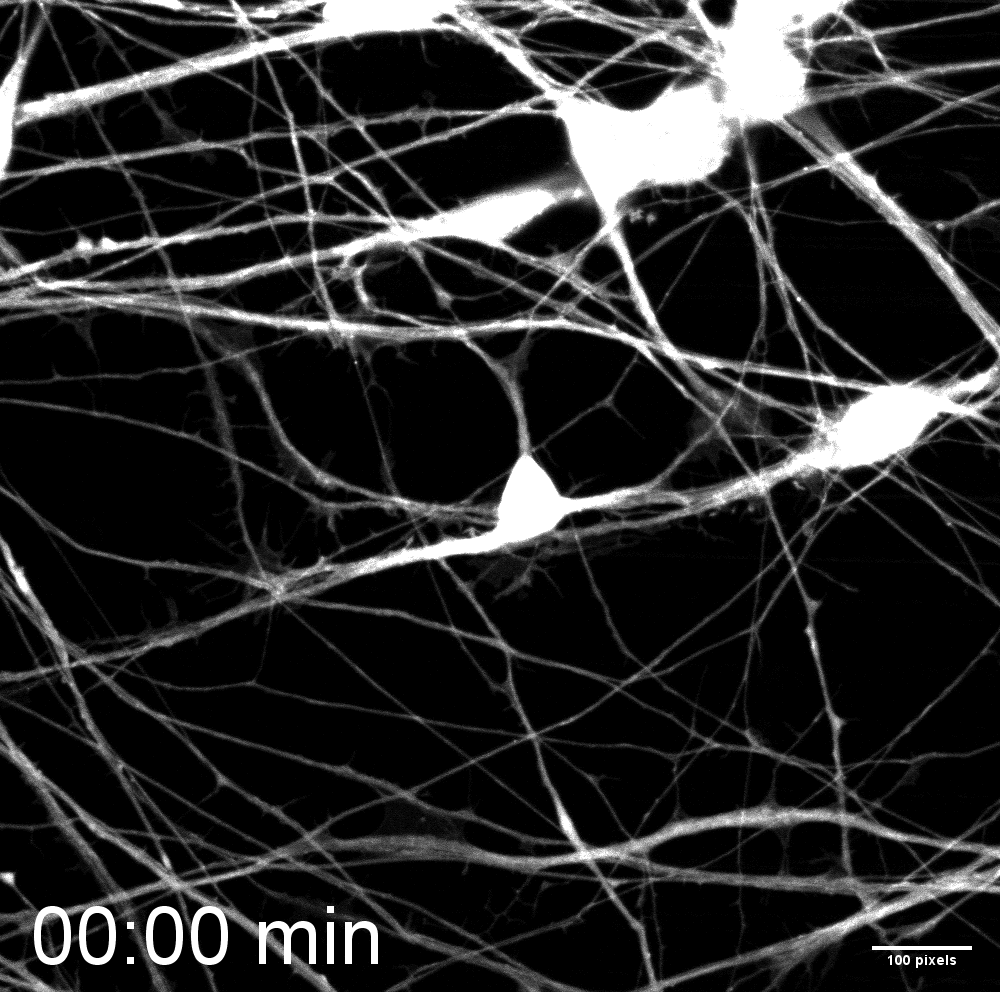

### S5 video

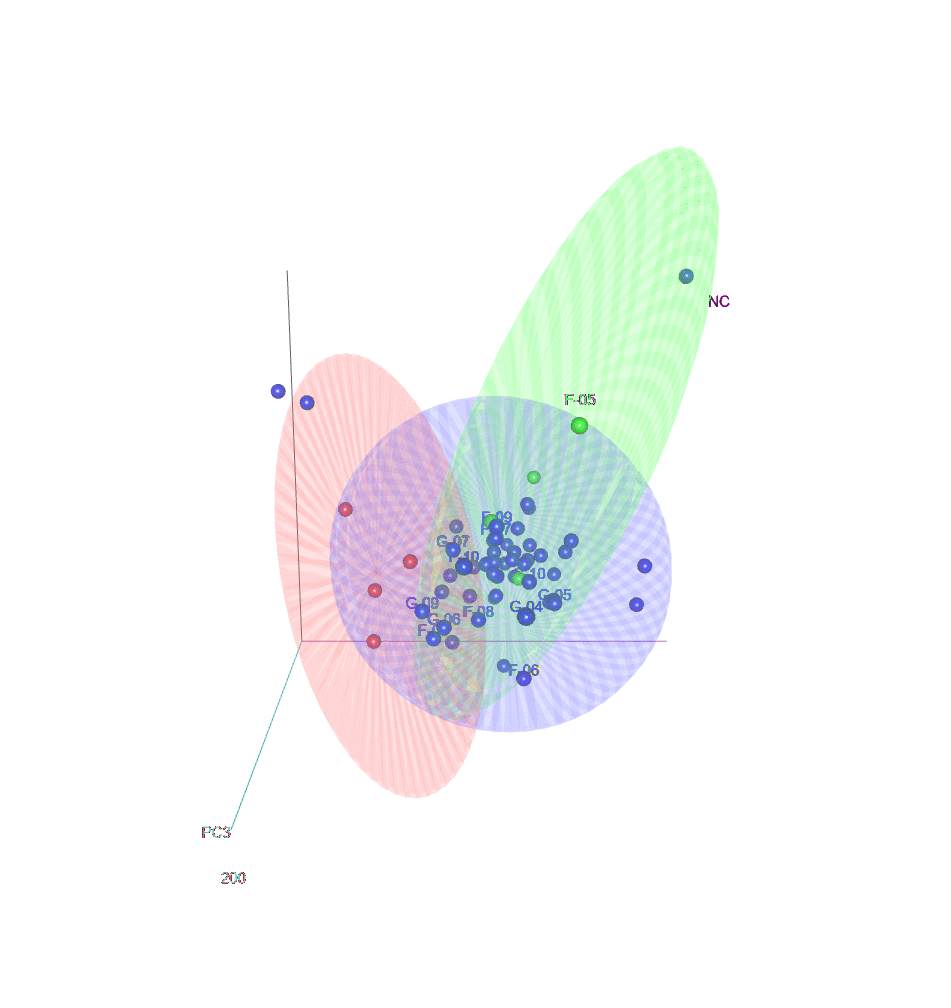

### S6 video

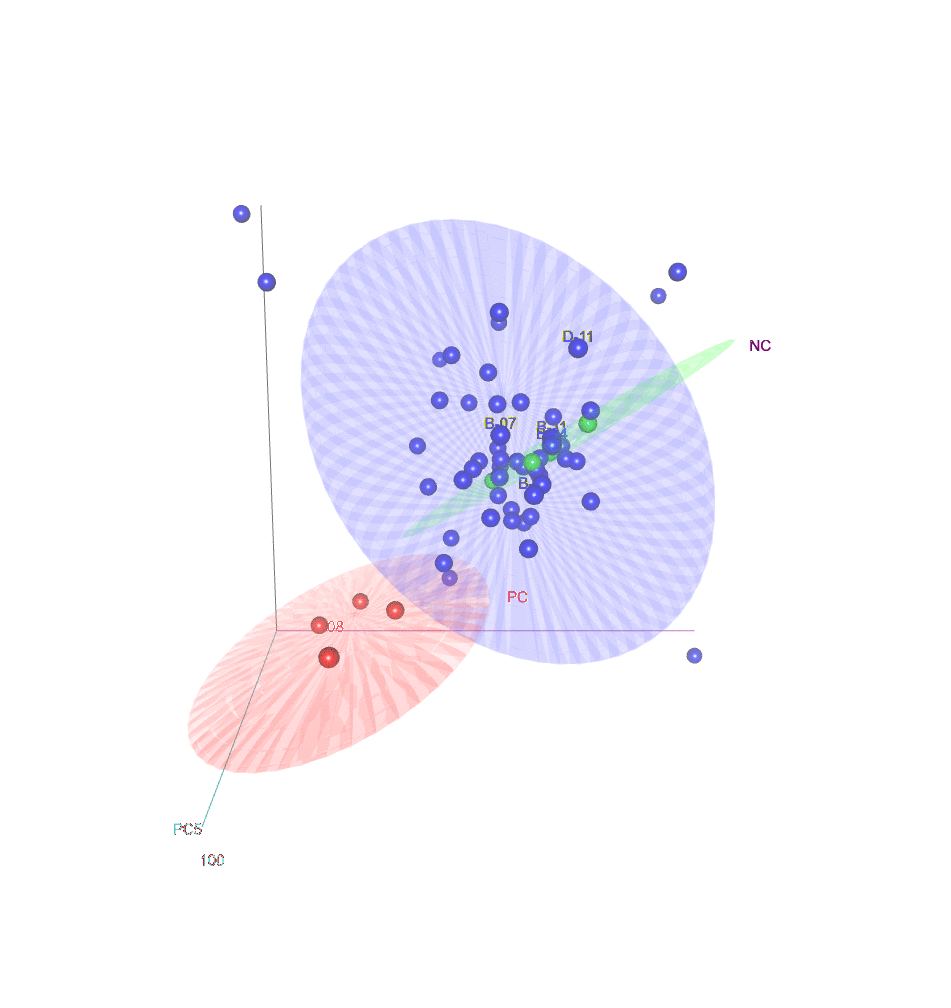

### S7 video

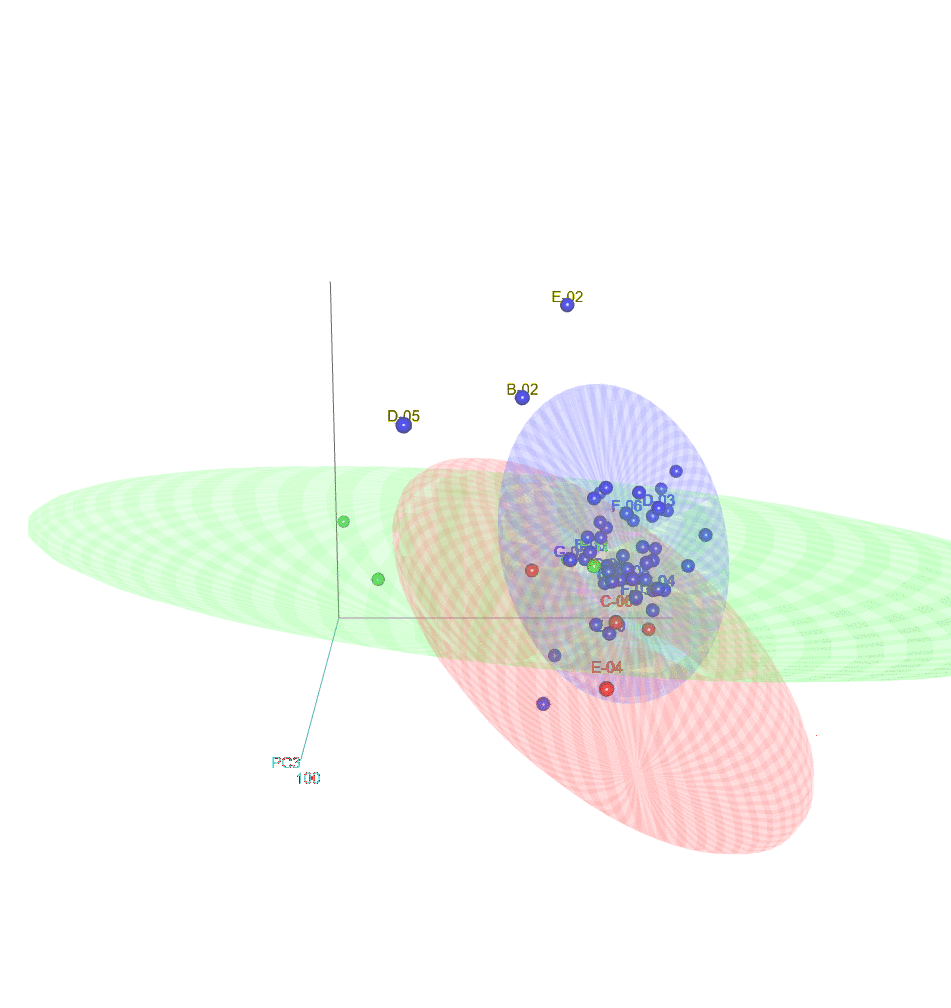

### S8 video

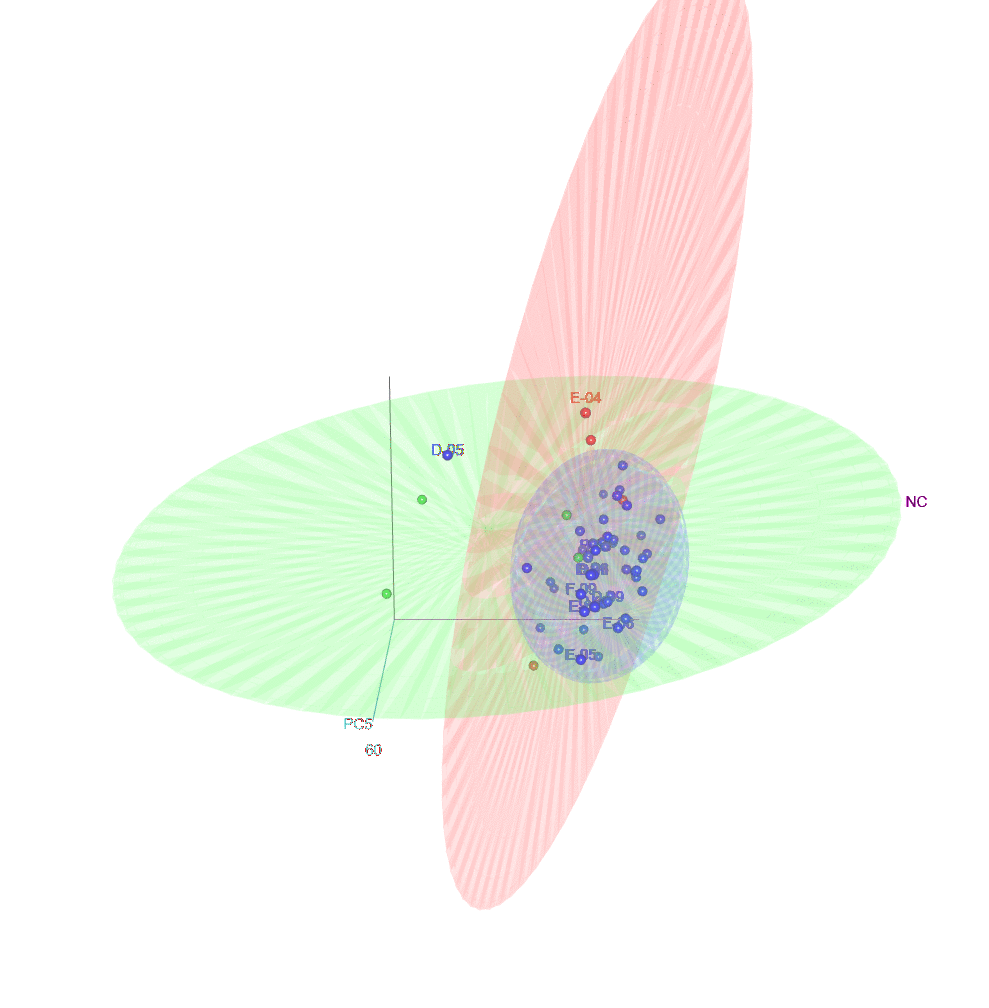

### S9 video

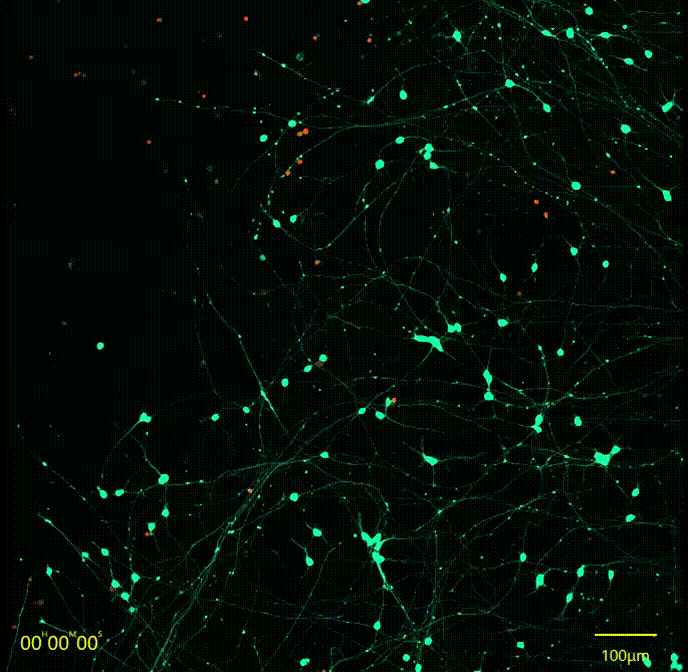

### S10 video

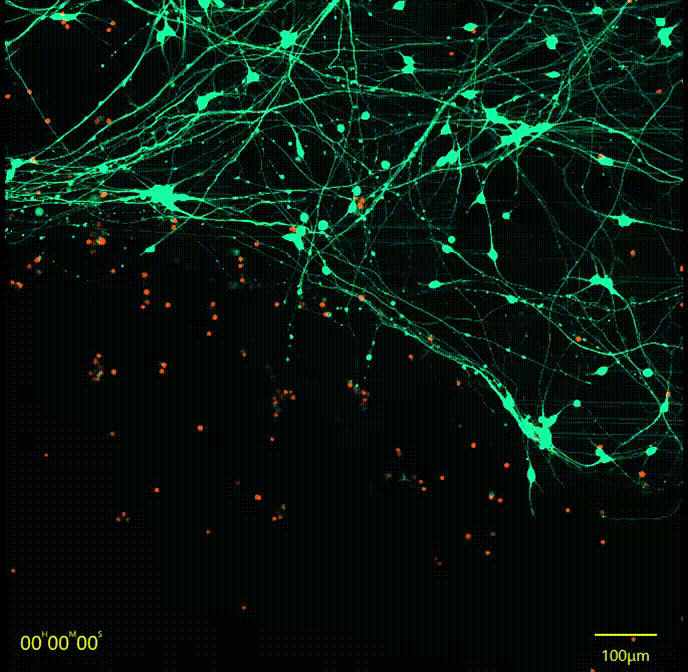

### S11 video

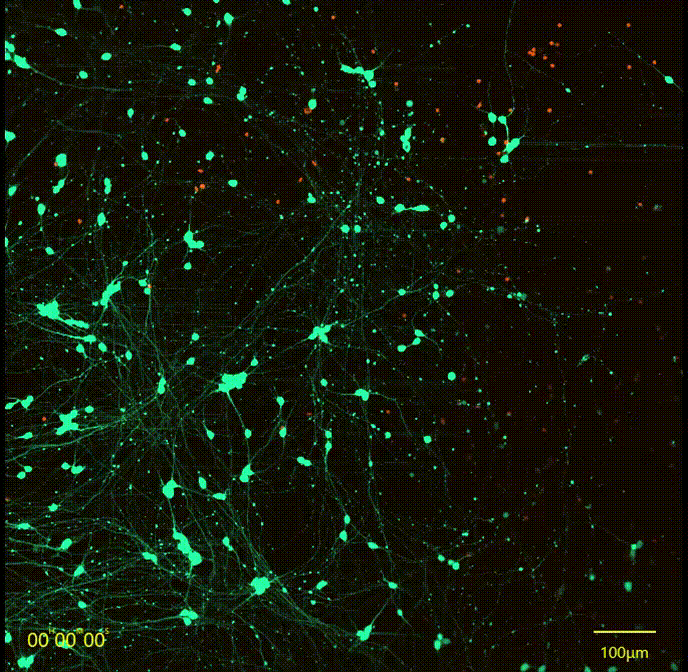

### S12 video

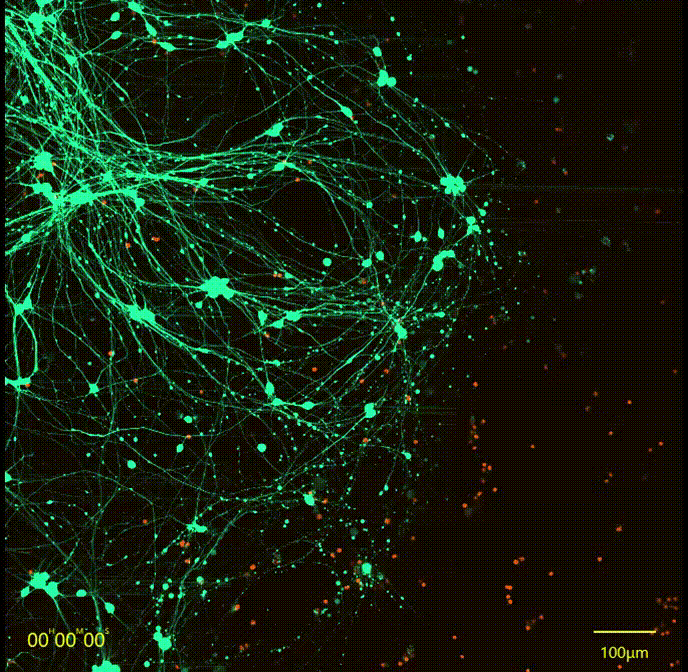

### S13 video

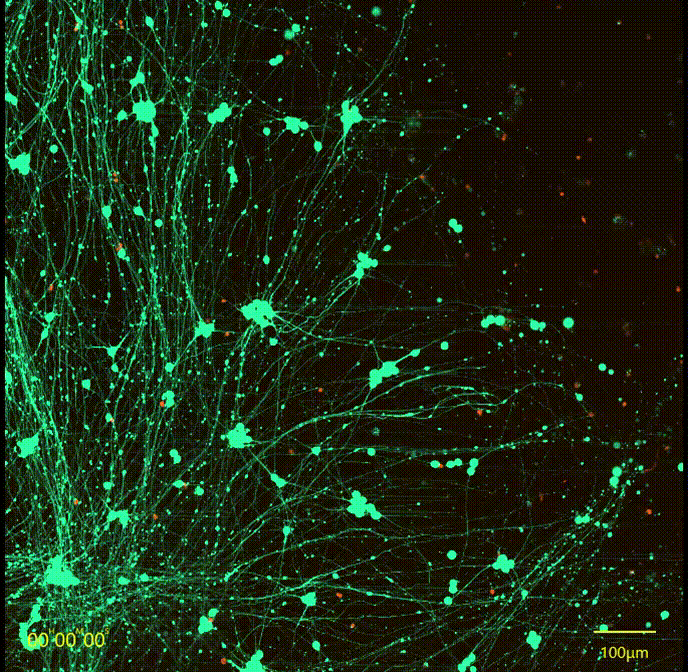

### S14 video

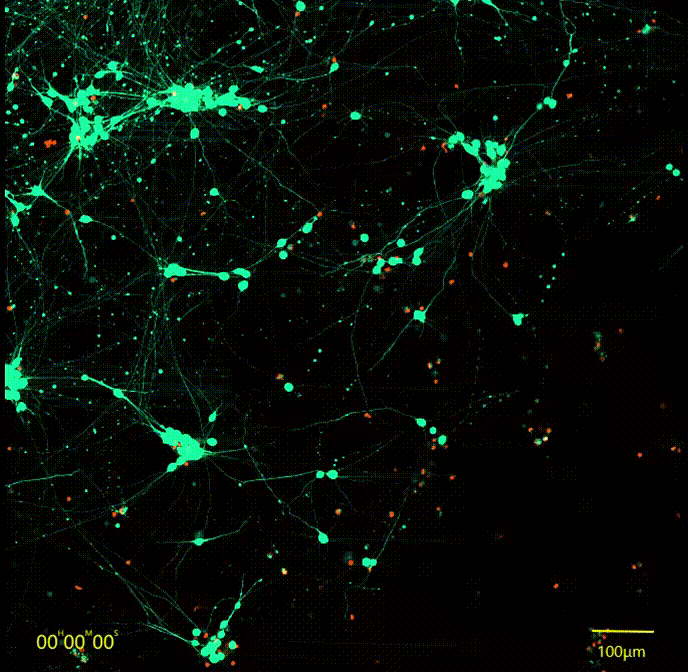
